# Faf2/Ubx2 potentiates p97/Cdc48 segregase activity via functional priming and reinforcement of the UT3 domain in Ufd1

**DOI:** 10.64898/2026.02.26.708172

**Authors:** Yuxiang Ren, Qingyun Zheng, Yujuan Duan, Zhengyang Xu, Qian Qu, Yicheng Weng, Siyi Cui, Yuanyun Yu, Man Pan, Lei Liu

**Affiliations:** The International Peace Maternity and Child Health Hospital, School of Medicine; Institute of Translational Medicine, National Center for Translational Medicine; School of Chemistry and Chemical Engineering; Shanghai Key Laboratory for Antibody-Drug Conjugates with Innovative Target, Shanghai Jiao Tong University, Shanghai, China; State Key Laboratory of Membrane Biology, Beijing Frontier Research Center for Biological Structure, School of Life Sciences, Tsinghua University; Tsinghua-Peking Joint Center for Life Sciences, Beijing, China; New Cornerstone Science Laboratory, Tsinghua-Peking Joint Center for Life Sciences, Ministry of Education Key Laboratory of Bioorganic Phosphorus Chemistry and Chemical Biology, Center for Synthetic and Systems Biology, Department of Chemistry, Tsinghua University, Beijing, China; School of Pharmaceutical Science, Shanghai Key Laboratory for Antibody-Drug Conjugates with Inno-vative Target, Shanghai Jiao Tong University, Shanghai, China

**Keywords:** Ubiquitin, p97/Cdc48, Faf2/Ubx2, boosted segregase activity, enzymatic ubiquitinated substrate synthesis

## Abstract

The AAA+ segregase p97/Cdc48 extracts poly-ubiquitinated proteins from entrenched cellular environments to maintain proteostasis. However, its functional capacity is not uniform; extracting substrates from dense assemblies like stress granules or membrane necessitates a fully potentiated state. Here, using precisely synthesized ubiquitinated substrates, we show that the Faf2/Ubx2 cofactor hyperactivates p97/Cdc48, lowering its minimal ubiquitin chain requirement and broadening linkage specificity. Cryo-EM analysis of the activated complex demonstrates that Faf2/Ubx2 potentiates p97/Cdc48 segregase activity by structurally remodeling the substrate recognition module (K48-diUb^Prox^) and priming its engagement with the AAA+ motor (p97/Cdc48-Npl4). These coordinated actions establish a ‘pump-unit’ architecture that drives efficient substrate processing. Our findings illuminate a conserved mechanism that unlocks the maximal capacity of p97/Cdc48 and provide a blueprint for therapeutically targeting its hyper-activated state in disease.

## Introduction

The AAA+ ATPase p97, also known as valosin-containing protein (VCP) or Cdc48 in S. cerevisiae, is a central orchestrator of cellular proteostasis^1–3^. It performs the essential mechanical work of extracting polyubiquitinated proteins from entrenched cellular environments—including membranes, organelles, chromatin, and stress granules—before unfolding them for degradation by the 26S proteasome^4–6^. This activity underpins critical processes from endoplasmic reticulum-associated degradation (ERAD) to epigenetic regulation, directly linking p97 dysfunction to neurodegenerative diseases and positioning it as a promising therapeutic target in oncology and virology^7–9^.

The indispensable function of p97/Cdc48 in proteostasis has driven sustained efforts to delineate the precise mechanisms by which it processes its substrates^10–23^. While p97/Cdc48’s fundamental role in processing ubiquitinated clients is well established, emerging evidence reveals that its functional capacity is not uniform, but rather exists in distinct activity states that govern its efficacy in substrate dislocation and unfolding. Critically, p97/Cdc48 in complex with canonical cofactors such as Ufd1-Npl4 (UN), often exhibits limited efficiency in extracting membrane-integrated ubiquitinated proteins, pointing to a “basal” activity state^24,25^. Very recent advances uncover the human cofactor Faf2 (Ubx2 in *S. cerevisiae*) enhances the system by recruiting and further activating the p97-UN complex^24,26^. This promotes the efficient extraction of recalcitrant substrates like ubiquitinated G3BP1 from stress granules, ultimately leading to rapid granule disassembly^27^. Furthermore, this Faf2-potentiated, ubiquitination-dependent, p97 activity extends to other proteostasis challenges, as demonstrated by its role in suppressing the aggregation of tau protein^28^.

Given the strong association between p97 dysfunction and neurodegenerative diseases as well as cancer, deciphering the molecular logic of its state-dependent activation is pivotal for developing targeted therapeutics. Such strategies aim to selectively modulate pathogenic p97 activities while sparing its essential global proteostatic functions. Thus, mechanistic dissection of p97’s functional states is imperative. While recent structural and biochemical insights have delineated the mechanism underlying the “basal” activity state of the Cdc48-UN segregase^10–23^, a fundamental gap persists: how does the cofactor Faf2 override or synergize with this canonical UN-mediated pathway to potentiate p97/Cdc48’s activity? Resolving this critical barrier is essential for a complete understanding of p97’s proteostatic repertoire and its therapeutic targeting.

Through an integrated approach combining protein engineering, biochemical characterization, cryo-EM structural analysis, AI-assisted modeling, and molecular dynamics simulations, we here identified a Faf2/Ubx2-drived hyperactivated state of the p97/Cdc48 segregase and uncovered the first structural characterization of Ufd1’s elusive UT3 domain. Specifically, Faf2 binds the UT3 domain of Ufd1, strengthens its proximal K48-linked diUb-binding activity, and positions it directly above the unfolded-Ub-binding groove of Npl4, thereby enabling efficient processing of polyubiquitinated substrates (including even branched Ub chains) and polypeptide translocation through the central pore of Cdc48/p97. Collectively, our results establish a model in which p97/Cdc48-dependent quality control systems achieve protein extraction and segregation by operating at full segregase capacity.

## Results

### Faf2 potentiates the unfolding activity of the p97 segregase and confers broadened tolerance to branched K48-polyubiquitin chains

Faf2 is composed of four major regions: a ubiquitin-associated (UBA) domain for recognizing polyubiquitinated substrates, a UAS thioredoxin-like domain responsible for binding unsaturated long-chain fatty acids, a ubiquitin-binding X (UBX) domain for p97/Cdc48 binding and a hydrophobic membrane anchor (MA) region that targets Faf2 to membrane organelles (**Figure 1A**)^27,29,30^. To directly assess Faf2’s functional impact, we generated a soluble full-length Faf2 construct by replacing its MA domain with a (GS)□□ linker (**Figure S1A** and **S1B**). Using a previously established substrate comprising N-terminally tagged mEos3.2 (Eos) conjugated to a K48-polyUb chain (>15 Ub units) (**Figure S1C**)^13,18,22^, we monitored the substrate unfolding via the irreversible loss of Eos fluorescence. We found that the addition of Faf2 significantly accelerated substrate unfolding (**Figure 1B**), increasing the relative initial rate by approximately 2.7-fold (**Figure 1C**).

**Figure 1.**
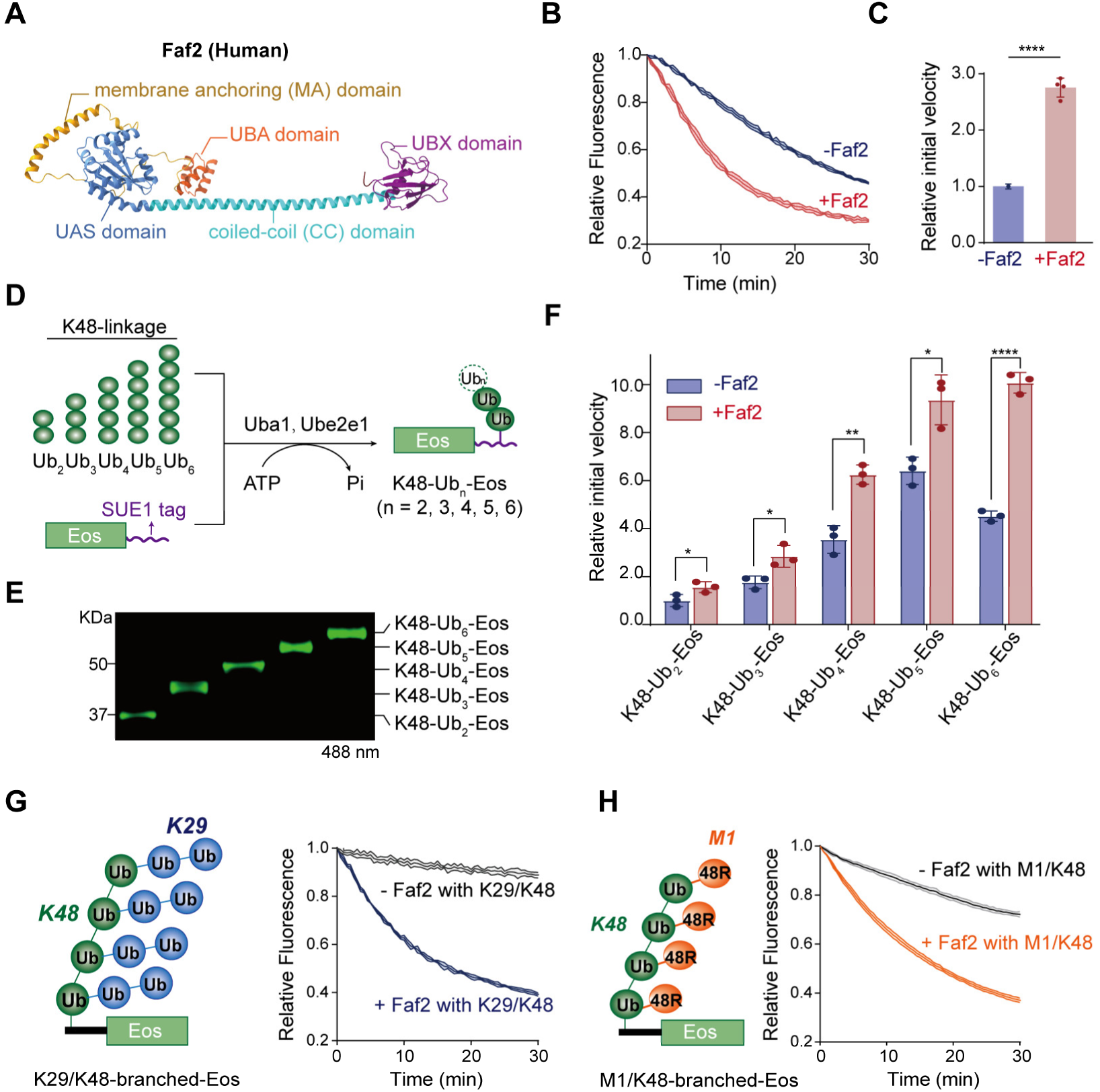
Faf2 accelerates p97-Ufd1-Npl4-mediated substrate unfolding and exhibits broad tolerance to branched K48-polyubiquitin chains. (**A**) AlphaFold3-predicted structural model of human Faf2. The membrane anchoring domain (MA) is shown in yellow, the ubiquitin-associated domain (UBA) in orange, the UAS domain in blue, the coiled-coil region (CC) in cyan, and the C-terminal UBX domain in purple. (**B**) The unfolding activity of p97-Ufd1-Npl4 towards K48-polyUb-Eos in the absence or presence of Faf2. The relative fluorescence signal was monitored for 30 min. (**C**) Quantification of initial velocities from (B). Initial rates were determined by linearly regression of early the initial phase and normalized against the condition without Faf2 (-Faf2). Error bars represent mean ± s.d. (n = 4 independent experiments). ****, *p* < 0.0001 (two-tailed Student’s *t*-test). (**D**) Schematic illustration of the enzymatic synthesis of K48-Ub_n_-Eos. Pre-assembled K48-Ub_n_ of varying lengths (Ub_2_ – Ub_6_) were conjugated to the SUE1 tag of Eos in the presence of Uba1 (E1), Ube2e1 (E2), and ATP. (**E**) SDS-PAGE analysis of purified K48-Ub_n_-Eos substrates with defined ubiquitin chain length (n = 2 – 6), visualized by fluorescence scanning at 488 nm. (**F**) Initial unfolding velocities of K48-Ub_n_-Eos substrates (n = 2 – 6) were measured in the absence (blue) or presence (red) of Faf2. Data are plotted as the ratio relative to the initial velocity of K48-Ub_2_-Eos without Faf2. Error bars represent mean ± s.d. (n = 3 independent experiments). Statistical significance was determined by two-way ANOVA. *, *p* < 0.05, **, *p* < 0.01, ****, *p* < 0.0001. (**G**) Unfolding of K29/K48-branched Ub chains. Left: Schematic of the K29/K48-branched-Eos substrate, consisting of K29-linked ubiquitin chains (blue) branched from a K48-linked backbone (green). Right: Unfolding kinetics of K29/K48-branched-Eos by p97-UN in the absence (black) or presence (blue) of Faf2. (**H**) Unfolding of M1/K48-branched substrates. Left: Schematic of the M1/K48-branched-Eos substrate. The K48-linked backbone (green) is decorated with single ubiquitin moieties (orange, M1 linkage). Right: Unfolding kinetics of M1/K48-branched-Eos by p97-UN in the absence (black) or presence (orange) of Faf2.

We next investigated whether Faf2’s acceleration effect depended on the length of K48-polyUb chain. Employing the SUE1 strategy (sequence-dependent ubiquitination via UBE2E1)^31^, we transferred pre-isolated free K48-Ub□, Ub□, Ub□, Ub□, or Ub□ chains (**Figure S1D** and **S1E**) onto the C-terminal SUE1 tag of Eos, generating length-defined polyubiquitinated substrates ranging from K48-Ub□-Eos to K48-Ub□-Eos (**Figure 1D** and **1E**). Unfolding assays revealed that Faf2 significantly enhanced the unfolding efficiency for substrates bearing at least 4 Ub units (K48-Ub□-Eos), increasing the relative average initial rate by 1.8-fold (**Figure 1F** and **S1F**). This enhancement was maximal for the K48-Ub□-Eos substrate, where the relative initial rate was elevated by 2.2-fold (**Figure 1F** and **S1F**).

Beyond homogeneous K48-polyUb chain, recent studies have shown that various branched polyUb chains also promote substrate degradation^32–37^. However, our prior biochemical reconstitutions failed to detect significant stimulation by branched chains^31^, implying a requirement for additional cofactors. To test whether Faf2 fulfills this role, we constructed an Eos substrate modified with a canonical K29/48-branched polyUb chain (**Figure S2A**). Briefly, we first isolated a shorter K48-polyUb-Eos substrate (10-14 units), which was then treated with a specific enzymatic cascade (Uba1-Ubc4-Ufd4) identified previously that catalyzes K29-polyUb chain extension^38,39^, yielding K29/48-branched-Eos. Strikingly, the native p97-UN complex exhibited minimal segregase activity against this branched substrate. In contrast, Faf2 addition dramatically enhanced unfolding efficiency (**Figure 1G**), increasing the relative initial rate by 7.2-fold (**Figure S2B**).

We further confirmed Faf2’s role using a structurally defined noncanonical M1/48-branched-Eos substrate. This substrate was generated using linear M1-diUb (with a distal K48R mutation to prevent spurious linkages) as a donor for conjugation to degron-Eos, ensuring exclusive N-terminal extension within the K48-polyUb chain (**Figure S2C**). Using this well-defined substrate, we again observed a significant stimulatory effect of Faf2 (**Figure 1H** and **S2D**), confirming its broad relevance for branched chain processing.

Collectively, our biochemical results reveal that Faf2 not only enhances the p97 segregase-mediated unfolding of K48-polyUb substrates but also is critical for efficient unfolding of branched polyUb substrates, thereby expanding the linkage tolerance of the p97 segregase complex.

### Architecture of an Ubx2-activated substrate-engaged Cdc48 complex

Next, we aimed to elucidate the structural mechanism by which Faf2 enhances the substrate-processing capability of the p97 segregase complex. Given that previous cryo-EM structures of the human p97-UN complex (in the presence and absence of polyUb substrates) failed to resolve clear density for the central tower-like domain of Npl4 (a region crucial for binding K48-polyUb substrates)^18,16^, we turned to the *S. cerevisiae* orthologous system. We purified the corresponding *S. cerevisiae* proteins including Cdc48, yNpl4, yUfd1, and Ubx2. Biochemical reconstitution confirmed that Ubx2 also enhances the linkage tolerance of the Cdc48/Npl4/Ufd1 complex (**Figure S3A-S3D**).

To favor a substrate-engaged state, we introduced the E588Q mutation into Cdc48 as previously described^15^. The assembled complex, comprising Cdc48^E588Q^, yNpl4, yUfd1, Ubx2, and K48-polyUb-Eos, eluted as a single peak in size-exclusion chromatography (**Figure S3E** and **S3F**). To capture the ATP-bound working state, 2 mM ATP was added to the purified complex immediately prior to grid preparation for cryo-EM single-particle analysis. Three-dimensional (3D) classification and refinement yielded an overall density map at 3.1 Å resolution (**Figure S4** and **Table S1**). In this EM map, Ubx2 density is visible but insufficiently resolved for atomic modeling due to intrinsic flexibility, whereas the Cdc48-yNpl4 density is clearly resolved (**Figure S3G**). The yNpl4 tower is positioned atop the Cdc48 D1 ring. Three folded Ub moieties are bound at the tower’s apex, while one unfolded Ub is engaged within the yNpl4 groove (**Figure S3H-S3K**). This architecture is consistent with the unfolding-enhanced conformation reported for the Cdc48-yUfd1-yNpl4 complex bound to a SUMO-polyUb substrate^40^. Notably, the Cdc48 D2 ring is completely invisible even after extensive unaligned 3D classification, these observations indicate a fundamentally distinct mechanism of Ubx2-mediated Cdc48 activation compared to that induced by SUMO-polyUb substrates.

To capture a more stable conformation of the Ubx2-activated, substrate-engaged Cdc48-UN complex, we employed two additional key strategies: replacing K48-polyUb-Eos with a length-defined free K48-Ub□ chain, and introducing an A494F mutation in yNpl4 to enhance K48-polyUb binding (**Figure 2A** and **2B)**. Applying analogous procedures to this new complex, we performed single-particle cryo-EM analysis and obtained a map with an overall resolution of 3.2 Å (**Figure S5A, S5B** and **Table S1**). Consistent with above polyUb-Eos-engaged Cdc48 structures, the D2 ring of Cdc48 remains invisible, and the Npl4 tower domain is positioned above the Cdc48 D1 ring, engaging two folded Ub units and one unfolded Ub (**Figure 2C** and **2D**). High-resolution features in these densities allowed for the unambiguous rigid-body docking of the known Cdc48-yNpl4 atomic structure (PDB: 6OA9).

**Figure 2.**
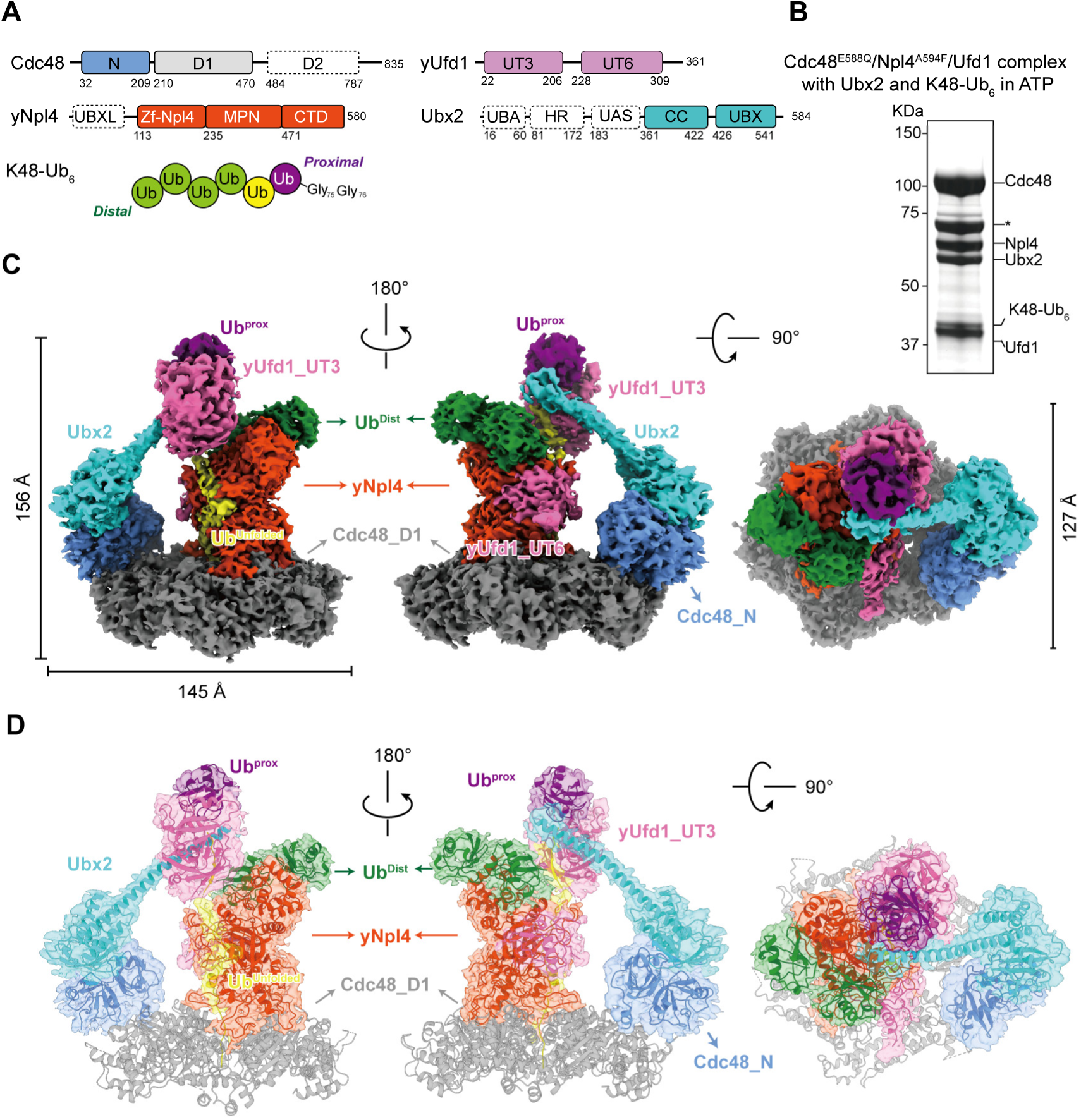
The structure of the Cdc48-yUfd1-yNpl4-Ubx2 complex with K48-Ub_6_. (**A**) Domain organization of Cdc48, yUfd1, yNpl4, Ubx2 and cartoon depiction of the K48-Ub_6_. Residue numbers at domain boundaries are indicated. The dashed box represents unresolved region. This color code for individual component is used throughout the article whenever possible. (**B**) SDS-PAGE analysis of the purified Cdc48-yUfd1-yNpl4-Ubx2-K48-Ub_6_ complex. The protein bands of the complex components are labeled as indicated. “*” represents Hsp70 protein. (**C and D**) Cryo-EM reconstruction (**C**) and the atomic model (**D**) of the complex in three different orientations.

Notably, our strategy to stabilize the complex markedly improved Ubx2 density quality. Further mask-based skip-alignment 3D classification and local refinement increased the local resolution to ∼4.1 Å (**Figure S5C**), permitting clear visualization of the Ubx2_CC-UBX and yUfd1_UT3 domains as well as a single Ub. The map reveals that yUfd1_UT3 is bound to the Ubx2_CC domain and the Ub. Moreover, based on the known function of the yUfd1_UT3 domain in binding the proximal Ub (Ub^prox^, purple) and facilitating unfolding of the adjacent donor Ub by Npl4^20^, the density surrounding UT3 may also encompass the flexible C-terminal tails of the unfolded Ub (Ub^unfolded^, yellow). Guided by AlphaFold3 structure predictions^41^, we constructed model of the Ubx2_CC-UBX/Ub^prox^/yUfd1_UT3/Ub^unfolded^/Cdc48_N complex, which was unambiguously as rigid body docked into the local map (**Figure 2D** and **3A**).

**Figure 3.**
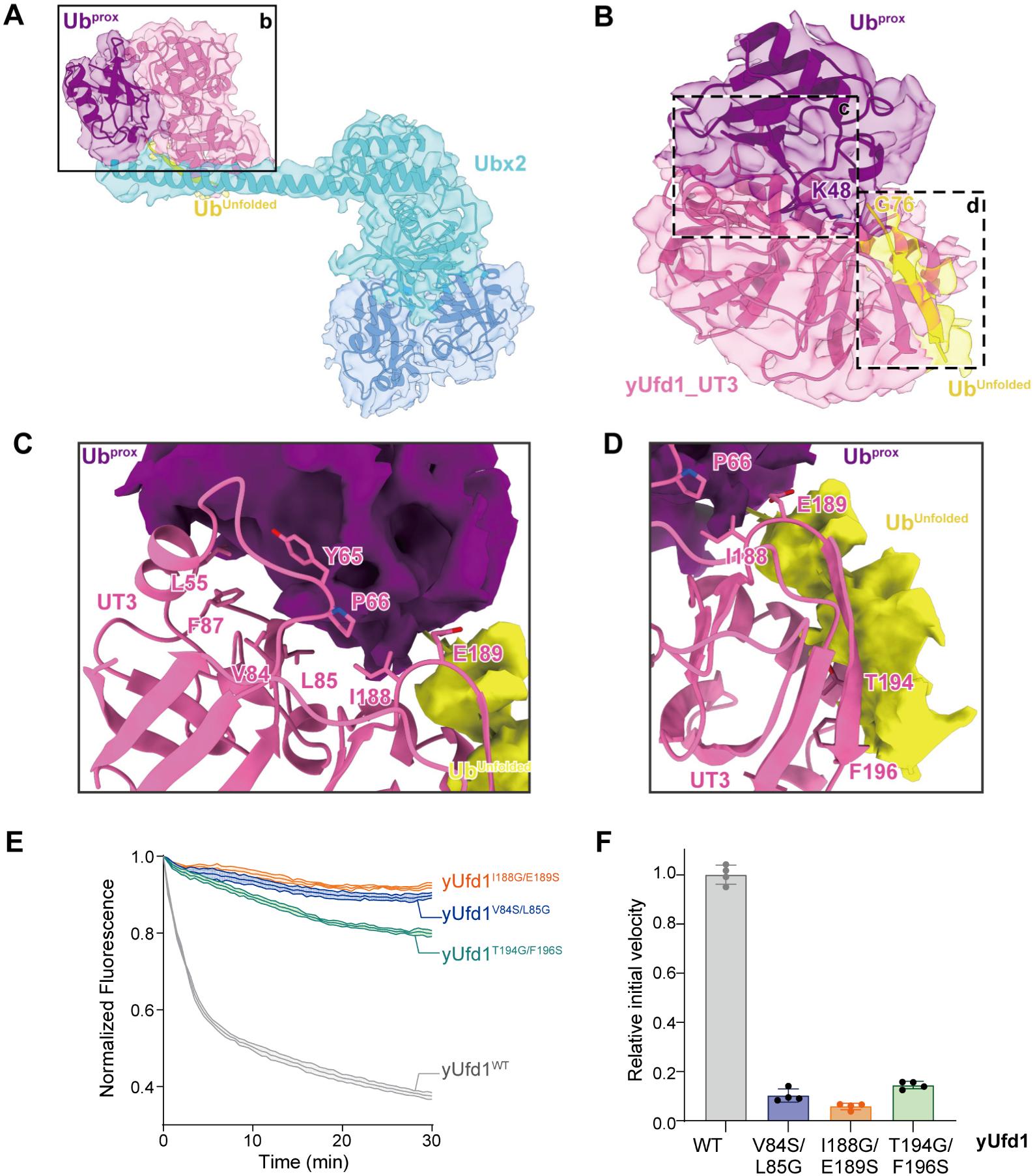
Integrative structural model of the Cdc48^NTD^-Ubx2^CC-UBX^-yUfd1^UT3^-Ub^prox^ and C-terminus of unfolded ubiquitin (Ub^unfolded^) complex. (**A**) An integrative model constructed by docking AlphaFold 3-predicted coordinates into the experimental Cryo-EM density. The semi-transparent cyan surface represents the Cryo-EM density map of Ubx2, the ribbon structures depict the fitted atomic models of Ubx2_CC-UBX region (cyan), the yUfd1_UT3 domain (pink), and the K48-linked Ub^prox^-Ub^unfolded^ (purple/yellow). The density envelope confirms the spatial organization of the complex. (**B**) Detailed view of the interaction between yUfd1_UT3 (cartoon) and the diUb segments (surface representation), extracted from the integrative model. Dashed boxes (c and d) indicate two distinct Ub-binding interfaces. (**C**) Close-up of box c, highlighting key residues on yUfd1_UT3 (e.g., V84, L85) that interact with the proximal ubiquitin (Ub^prox^, purple). (**D**) Close-up of box d, highlighting residues on yUfd1_UT3 (e.g., I188, E189, T194, F196) that engage the ubiquitin moiety positioned for unfolding (Ub^unfolded^, yellow). (**E**) The unfolding activity of Cdc48-yUfd1-yNpl4 towards K48-polyUb-Eos in the presence of different yUfd1 variants. The curves show the mean and standard deviation of four replicates. (**F**) Experiments as in (E) were done with selected yUfd1 mutants and controls. The initial unfolding rates were determined and normalized to that of WT. Shown are the mean and standard deviation of four replicates.

The final merged structure reveals Ubx2 adopting a conformation analogous to a “pumping unit”, defined by two principal interfaces: the UBX domain binding to the Cdc48_N domain, and the CC domain engaging both the yUfd1_UT3 domain and the proximal Ub of the substrate. This organization places the UT3 domain directly atop the yNpl4 tower (**Figure 2D**). Critically, these two features are linked by the unfolded Ub, with its N-terminus bound in the Npl4 groove and inserted into the Cdc48 central pore, and its C-terminus anchored at UT3, thereby establishing a critical spatial connection.

### Visualization of the interaction between the UT3 domain of Ufd1 and K48-linked polyubiquitin

Focusing on the head of the “pump unit”, we captured the first visualization of the UT3 domain’s interaction with K48 diUb. The UT3 domain simultaneously contacts the Ub^prox^ and the Ub^unfolded^ (**Figure 3B**). Critically, UT3 engages Ub^prox^’s canonical patches (centered on I44 and L8) via a large hydrophobic region comprising L55, F87, V84, L85, and I188, supplemented by peripheral polar interactions (**Figure 3C** and **3D**). This binding interface extends beyond the previously characterized interaction, which was restricted to UT3’s L55 and Ub^prox^’s I44 (**Figure S6A**)^20^. Mutation of key UT3-Ub^prox^ interface residues (V84S/L85G and I188G/E189S) severely impaired the unfolding activity of the Cdc48-UN complex (**Figure 3E** and **3F**), confirming UT3’s essential role as a Ub^prox^ sensor for directional ubiquitinated substrate anchoring.

For the interaction between Ub^unfolded^ and UT3, T194 and F194 of UT3 form a network of backbone hydrogen bonds with L71 and L69 of the Ub^unfolded^ C-terminus (**Figure 3D** and **S6B**). This binding mode stabilizes the flexible Ub C-terminal tail of the Ub^unfolded^ and locks it onto the UT3 surface. Mutation of key UT3-Ub^unfolded^ interface residues (T194G/F196S) severely impaired the unfolding activity of the Cdc48-UN complex (**Figure 3E** and **3F**).

### Functional investigation into CC-UBX domain of Faf2/Ubx2

We next analyzed Ubx2 within the activated substrate-engaged Cdc48 complex. The UBX domain of Ubx2 engages the Cdc48_N domain through a conserved recognition mode (**Figure S6C**), whereas the Ubx2_CC domain bridges the “pump head” (UT3&Ub) to the “pump tail” (UBX & p97_N domain). Notably, a truncated version of Ubx2 containing only the UBX and CC domains retains full activity in Cdc48 unfolding assays, confirming that these two domains are sufficient to mediate this function (**Figure S6D**). Homology comparison confirmed the evolutionary conservation of the UBX and CC domain (**Figure 4A**). Consistent with observations in yeast, a minimal truncation of human Faf2 demonstrated that its CC-UBX module is also sufficient to activate the p97-UN complex (**Figure 4B** and **4C**). Structural analysis revealed that CC domain forms a broad interface (441.9 Å²) with UT3 through multivalent interactions (**Figure 4D**). Mutations disrupting the evolutionarily conserved residues (K372, Y379, S382, L383) at the homologous Faf2_CC interface (Faf2^CC-4mut^) severely impaired substrate unfolding activity (**Figure 4E** and **4F**).

**Figure 4.**
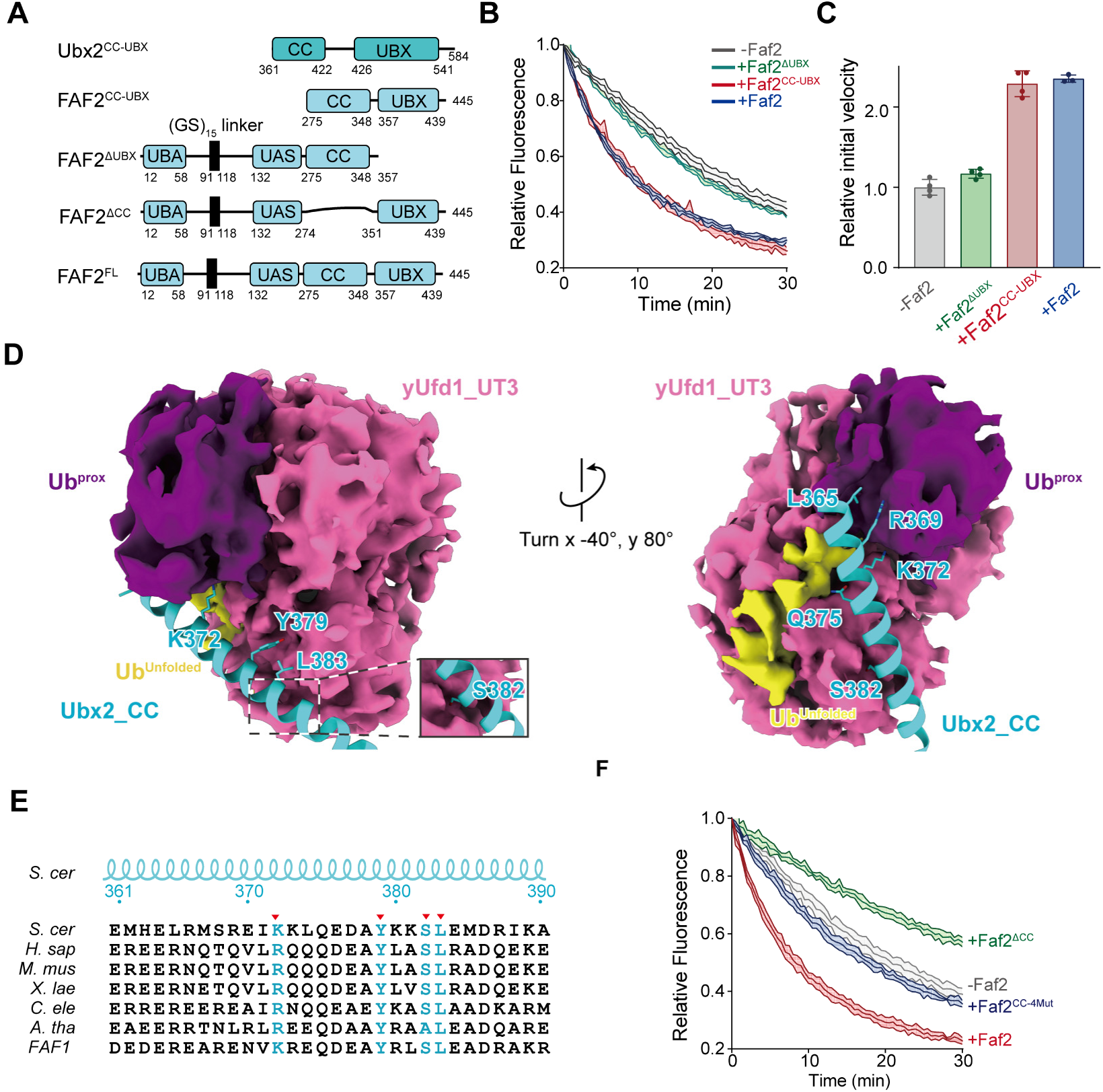
The CC domain is essential for Faf2/Ubx2-mediated stimulation of p97/Cdc48 activity. (**A**) Diagram of Faf2, Faf2 truncations lacking UBX (ΔUBX), coiled-coil region (ΔCC), or fragment of Faf2 and Ubx2 encompassing the coiled-coil region and UBX domain (CC-UBX). (**B**) Unfolding of K48-polyUb-Eos by p97-UN or p97-UN complexed with Faf2, Faf2^ΔUBX^ or Faf2^CC-UBX^. (**C**) Quantification of initial velocities from (B). Initial rates were determined by linearly regression of early the initial phase and normalized against the condition without Faf2 (-Faf2). Error bars represent mean ± s.d. (n = 4 independent experiments). (**D**) Detailed views extracted from the integrative structural model, illustrating the positioning of the Ubx2 coiled-coil (Ubx2_CC, cyan) within the composite interface formed by yUfd1_UT3 (pink) and the K48-diubiquitin segment (Ub^prox^/purple, Ub^unfolded^/yellow). Left panel: The conserved residues selected for mutagenesis (K372, Y379, S382, L383) are shown projecting towards the UT3 interface, suggesting a direct role in stabilizing the ternary complex. Right panel: A rotated view highlighting additional interfacial residues and the extensive surface complementarity between the coiled-coil helix and the ubiquitin. (**E**) Multiple sequence alignment of the selected coiled-coil (CC) segment (corresponding to residues 361-390 of *S. cerevisiae* Ubx2) across representative species and the human paralog Faf1. The four conserved residues highlighted in cyan and indicated by red triangles (▾) denote the positions targeted for site-directed mutagenesis (Faf2^CC-4mut^) in functional activity assays. (**F**) Unfolding of K48-polyUb-Eos by p97-UN or p97-UN complexed with Faf2, Faf2Δ^CC^ or Faf2^CC-4mut^.

To dynamically monitor the impact of the CC-UBX domain on UT3 recruitment of ubiquitinated substrates, we developed a FRET-based proximity assay (**Figure 5A**). We employed BODIPY-FL as the FRET donor and Texas Red as the acceptor. A synthetic N-terminal degron labeled with BODIPY-FL was ubiquitinated to generate degron^BODIPY^-K48-Ub_n_ (**Figure S7A** and **S7B**). Separately, we introduced an 11-residue ybbR tag into the Ufd1 N-segment. This tag enabled enzymatic, site-specific labeling with a Texas Red-CoA conjugate catalyzed by Sfp phosphopantetheinyl transferase^42^, yielding Ufd1^Texas^^Red^ (**Figure S7A** and **S7B**). Assembly of the p97-UN complex containing Ufd1^Texas^^Red^ and its proximity to degron^BODIPY^-K48-Ub_n_ generated a FRET signal (**Figure S7C**). Titration with increasing p97-UN concentrations yielded an apparent binding affinity (app. K_d_) of 9.7 nM (**Figure 5B**). Critically, addition of either full-length Faf2 or its isolated CC-UBX domain robustly enhanced the interaction affinity by approximately two-fold, indicating that this cofactor potentiates substrate processing by strengthening UT3-mediated engagement with the ubiquitinated substrate.

**Figure 5.**
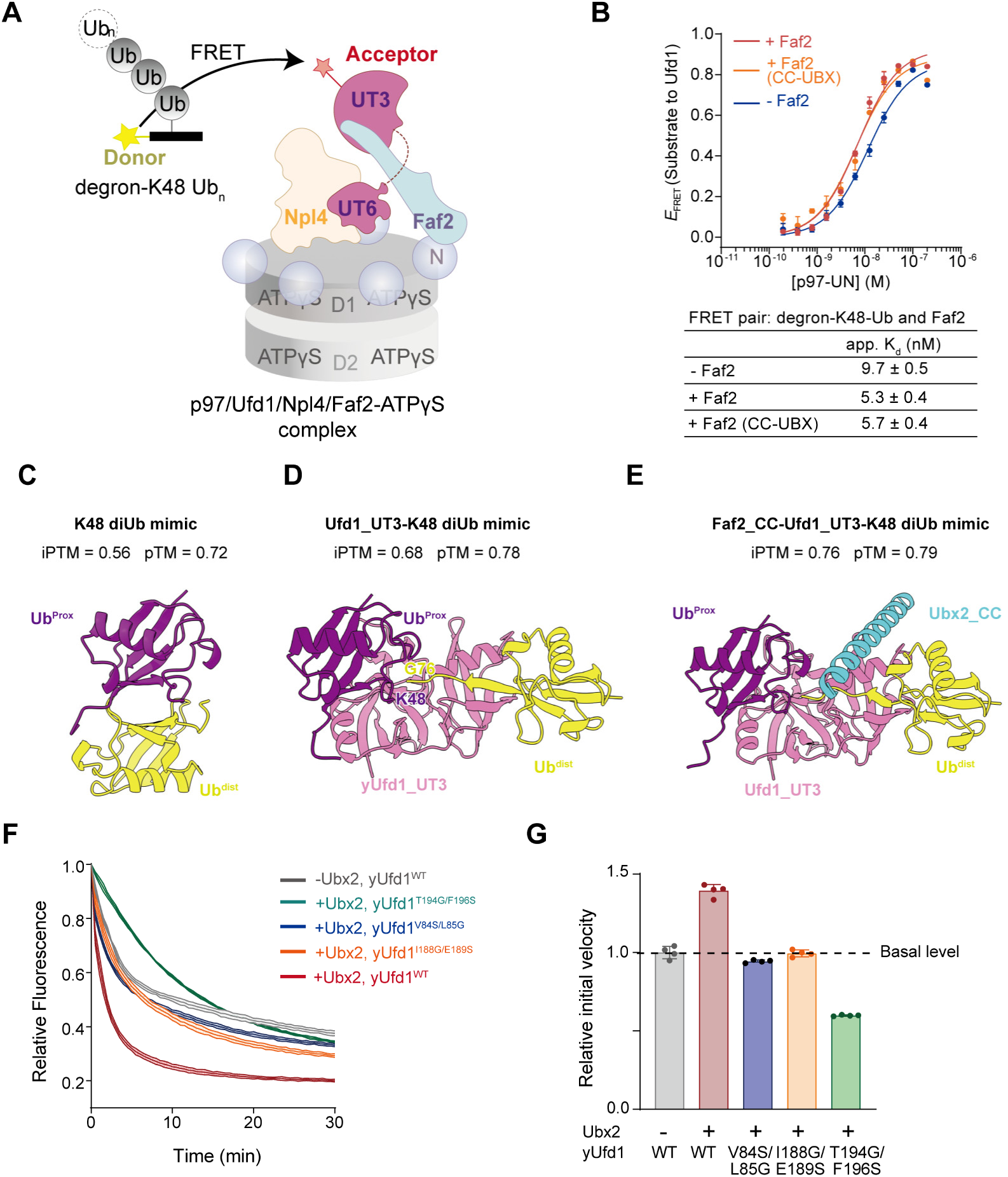
The Faf2/Ubx2 CC bridges Ufd1 and K48 diUb to reinforce substrate engagement. (**A**) Schematic representation of the FRET-based binding assay. The interaction between the donor-labeled substrate (degron-K48-Ub_n_) and the acceptor-labeled p97-UN complex was monitored in the presence of ATPγS. The acceptor fluorophore was attached to N-terminus of Ufd1-UT3 domain. (**B**) Equilibrium binding titrations of p97-UN and K48-linked polyubiquitinated substrate. Increasing concentrations of the Texas Red-labeled p97-UN complex were titrated against a fixed concentration of BODIPY-FL labeled degon-K48-Ub_n_ in the absence (blue) or presence of full-length Faf2 (red) or the Faf2^CC-UBX^ (orange). The FRET efficiency (*E*_FRET_) was plotted against the complex concentration to determine the apparent dissociation constant (app. *K*_d_). The table summarizes the app. *K*_d_ values (mean ± s.d., n = 4). Faf2 and its CC-UBX truncated form enhance the binding affinity to a similar extent. (**C-E**) AlphaFold 3 structural modeling of the K48-diUb mimic (**C**), the Ufd1_UT3-K48-diUb mimic (**D**) and the Faf2_CC-Ufd1_UT3-K48-diUb mimic (**E**), Predicted structural models and the corresponding confidence scores (iPTM and pTM) generated by AlphaFold 3 are shown. (**F**) The impact of Ubx2 on the substrate unfolding efficiency of Cdc48 complexes harboring various yUfd1 mutations. (**G**) Quantification of initial velocities from (F). Error bars represent mean ± s.d. (n = 4 independent experiments).

To elucidate the mechanism by which Faf2/Ubx2 accelerates engagement of ubiquitinated substrate, we employed a recently established AlphaFold3 modeling technique to study the polyubiquitin complex. We constructed a mimic of K48 diUb by mutating its linking residue to cysteine to form a disulfide bond, thereby mimicking the K48 isopeptide bond^43^. The predicted K48 diUb mimic revealed a characteristic “compact” conformation, consistent with previously reported structures (**Figure 5C**)^44^. Significantly, binding of the Ufd1_UT3 domain induced substantial extension and rotation of the distal Ub tail, transitioning the K48 diUb mimic into an “open” conformation (**Figure 5D**). The added Faf2_CC domain bound precisely within the space created between the extended Ub units (**Figure 5E**). Modeling results of the yeast system also indicated that Ubx2_CC adopts a conserved mechanism (**Figure S7D-7F**). We further calculated binding free energies the UT3-K48 diUb complex in the absence and presence of Faf2_CC using molecular mechanics/Poisson-Boltzmann surface area (MM/PBSA)^45–47^, and found that the binding of Faf2_CC significantly favored to the stability of UT3-K48 diUb complex (**Table S2**). These results demonstrate that the UT3 domain binds and triggers a conformational change in the K48-polyUb chain, with the Faf2/Ubx2_CC domain serving as a stabilizer of the resulting “open” conformation.

We therefore tested this hypothesis by asking whether Ubx2 could compensate for the loss of function caused by UT3 mutations. Remarkably, Ubx2 significantly rescued the activity of mutations at both the UT3-Ub^prox^ interface (e.g., V84S/L85G, I188G/E189S) and the UT3-Ub^unfolded^ interface (e.g., T194G/F196S) (**Figure 5F** and **5G**). This functional rescue confirms that Ubx2/Faf2 is crucial for supporting the UT3-mediated initial Ub engagement step. In addition, the observed rescues of p97 activity on K29/48-branched and M1/K48-branched Ub chains are consistent with our predicted complex structure. Adjacent Ub moieties conjugated to the proximal Ub likely induce potential Ub-Ub interactions at the UT3 binding interface. The CC domain counteracts these potential interactions by strengthening the UT3-K48 diUb interaction, thereby extending p97 segregase tolerance for K48-linked chains (**Figure S7G**).

## Discussion

In the previously characterized “basal” activity state of p97/Cdc48 complex, engagement of ubiquitinated substrates is initiated by an unfolded Ub moiety binding to Npl4’s groove-like surface, with its N-terminus inserting into the Cdc48 central channel^15^. Two subsequent folded Ubs then engage the Npl4 tower domain. The Cdc48 motor subsequently uses ATP hydrolysis to translocate the initiating Ub through the central channel, progressively unfolding the attached polyUb chain and ultimately threading the substrate protein for processing^20,21^. Biochemical analyses have indicated that the cofactor Ufd1, particularly its UT3 domain, provides critical specificity for K48-polyUb and is critical for the unfolding and processing of ubiquitinated substrates, yet the underlying molecular mechanism has remained elusive^20^.

Here our integrated study defines a Faf2/Ubx2-promoted hyperactive state that unlocks the full segregase potential of p97/Cdc48. We demonstrate that Faf2/Ubx2 functions not merely as a p97/Cdc48 adaptor but as a conformational catalyst: by binding the Ufd1 UT3 domain, it enhances UT3’s engagement with the proximal K48-diUb unit on the substrate. This interaction stabilizes an extended, “open” conformation of the K48-diUb, positioning it for optimal docking onto the Npl4 tower domain and mechanically priming it for processive translocation. Together, these findings establish a “pump unit” model wherein Faf2/Ubx2, via its CC-UBX module, bridges the substrate-recognition module (K48-diUb_UT3) to the p97/Cdc48_Npl4 ATPase motor, thereby dramatically enhancing processing efficiency and broadening linkage tolerance (**Figure 6A** and **6B**).

**Figure 6.**
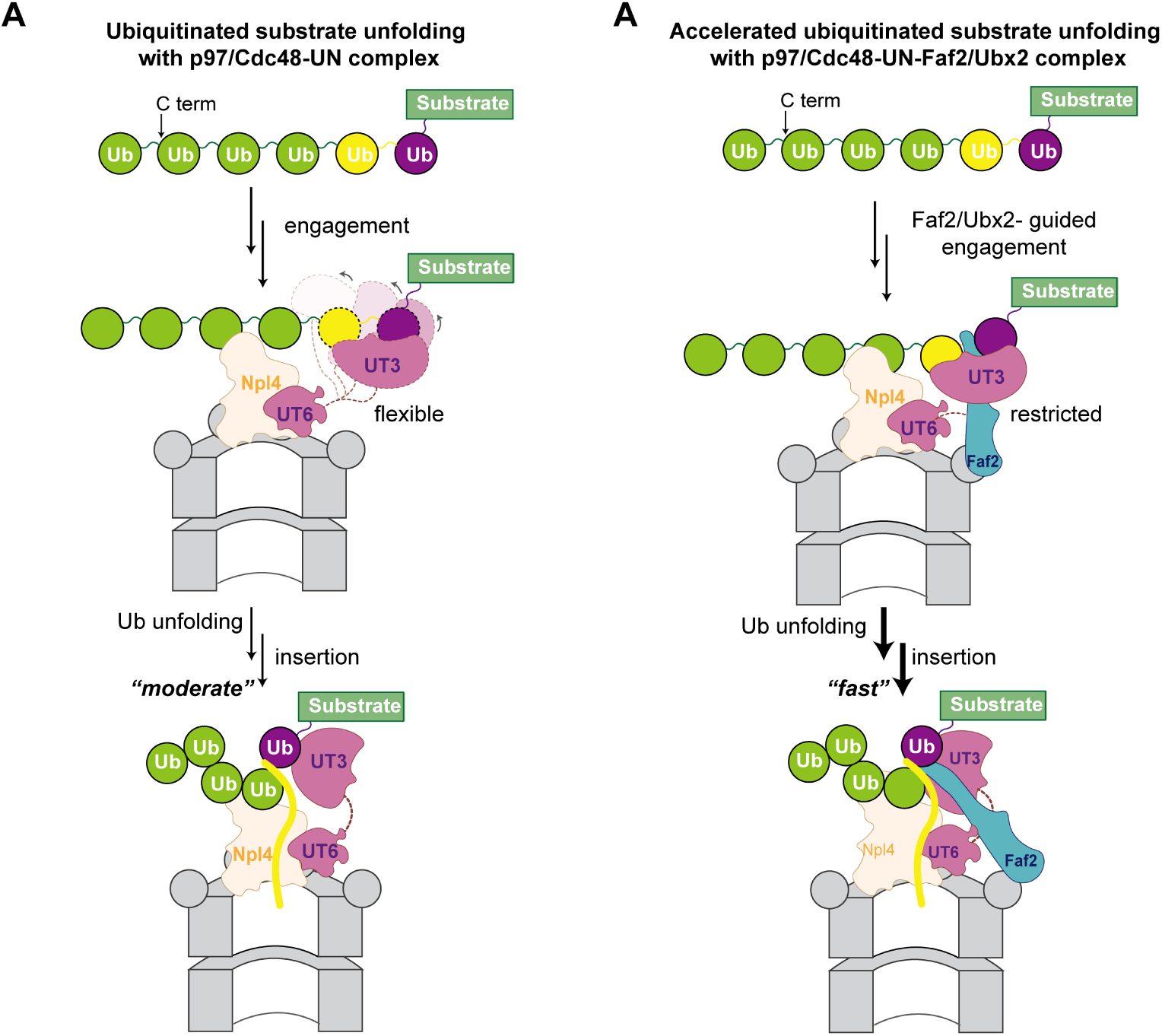
Model for accelerated p97/Cdc48 segregase activity by Faf2/Ubx2. (**A**) Model scheme of a fundamental p97/Cdc48 segregase activity. UT3 domain of Ufd1 is responsible for binding proximal K48 diUb, Npl4 is responsible for binding distal K48 diUb, UT3 continuously adjusts the dynamics with proximal K48 diUb to keep the Ub^unfolded^ unit for optimal engagement with the Npl4 tower and priming it for translocation through the p97/Cdc48 central pore. (**B**) Model scheme of a accelerated p97/Cdc48 segregase activity by Faf2/Ubx2. In the presence of Faf2/Ubx2, Faf2/Ubx2 binds the UT3 domain of Ufd1, strengthens its proximal K48-linked diUb-binding activity, and positions it directly above the unfolded-Ub-binding groove of Npl4, thereby enabling efficient processing of polyubiquitinated substrates and polypeptide translocation through the central pore of Cdc48/p97.

A central mechanistic insight from our work is the enhancement of UT3’s Ub binding affinity by the Faf2_CC domain. Our biochemical and AlphaFold3-guided analyses show that UT3 alone can induce an “open” chain conformation, but this state is inherently flexible. Faf2_CC acts as a molecular wedge and scaffold, binding within the space created between the extended Ubs and locking the UT3-Ub^prox^-Ub^unfolded^ assembly into a stable, high-affinity configuration. This explains the near two-fold increase in binding affinity and the robust functional rescue of UT3 interface mutants by yeast Faf2 homologue, Ubx2. Thus, Faf2 does not create a new binding site but rather potentiates a pre-existing, low-efficiency interaction within the canonical UN complex, converting it into a high-capacity substrate-processing unit.

Notably, the use of enzymatically synthesized ubiquitin chains with defined lengths and linkages proved instrumental in elucidating Faf2/Ubx2’s mechanism of p97/Cdc48 activation. These precision substrates demonstrated Faf2’s role in lowering p97’s minimal chain requirement (≥Ub_4_) and enabling branched chain processing—insights that were obscured when using heterogeneous ubiquitin preparations. And the K48-Ub□ chain played a pivotal role in stabilizing the Cdc48 complex, allowing high-resolution cryo-EM analysis of the activated complex. This work underscores how advances in ubiquitin ligation technologies continue to empower mechanistic studies of the ubiquitin-proteasome system.

This potentiation mechanism has profound implications for p97’s functional versatility in cells. Our biochemical data show that Faf2’s role becomes critical for substrates modified with shorter or branched ubiquitin chains. These complex topologies likely present greater steric or thermodynamic barriers to unfolding by the basal UN machinery. By stabilizing the UT3-mediated “open” conformation, Faf2 effectively lowers the activation energy for initial Ub remodeling and mitigates potential steric clashes that branched ubiquitination poses to proximal Ub engagement, enabling the processing of otherwise recalcitrant substrates. This explains why Faf2 is essential for the efficient disassembly of stress granules containing K63-linked ubiquitinated G3BP1^27^, a process requiring the extraction of proteins from dense, structured assemblies. Our model posits that Faf2 expands the operational range of p97, allowing it to handle a broader spectrum of ubiquitin signals and substrate complexities, thereby integrating specific cellular cues (e.g., fatty acids via its UAS domain) with global proteostatic demand^48^.

The evolutionary conservation of the CC-UBX module from yeast Ubx2 to human Faf2 underscores the fundamental nature of this activation mechanism. The finding that this minimal module is sufficient for full activity suggests it represents the core functional engine, with additional domains (UBA, UAS, MA) conferring contextual regulation, substrate targeting, and subcellular localization. This modular design allows p97’s activity to be precisely tuned by a family of UBXD proteins in different cellular compartments and pathways. Our work on Faf2 provides a paradigmatic framework for understanding how related cofactors might modulate p97 function in other contexts, such as in ERAD or DNA damage repair^49^.

In conclusion, we have delineated a molecular blueprint for p97/Cdc48 hyper-activation. This knowledge not only advances our fundamental understanding of AAA+ ATPase regulation but also opens new avenues for therapeutic intervention. Diseases driven by p97 dysfunction, such as inclusion body myopathy or neurodegenerative disorders, as well as cancers reliant on p97 for protein homeostasis, may be susceptible to modulators targeting the Faf2-UT3 interface or the induced “open” conformation of the Ub chain. Such strategies could allow precise tuning of p97 activity with greater specificity than targeting the essential ATPase core itself.

## Acknowledgments

We thank the National Natural Science Foundation of China (92253302, T2488301, 22137005, and 22227810 to L.L.; 22277073 and 92253302 to M.P.; 22407085 to Q.Z.), National Key R&D Program of China (2022YFC3401500 to L.L.; 2025YFA1310000, 2023YFA0915300 to M.P.), New Cornerstone Science Foundation (to L.L.), Shanghai Pilot Program for Basic Research – Shanghai Jiao Tong University (21TQ1400224 to M.P.), Fundamental Research Funds for the Central University (to M.P.), We thank for cryo-EM data collection in the Instrument Analysis Center (IAC), Shanghai Jiao Tong University on Cryo-EM. We also acknowledge the National Facility for Translational Medicine (Shanghai) for support.

## Author contribution

L.L, M.P., Q.Z. proposed the idea, designed the experiments and supervised the experiments. Y.R. and Q.Z. conducted the biochemical experiments. Y.R. prepared cryo-EM sample and FRET assays. Y.D. conducted the cryo-EM experiments. Z.X. conducted the Alphafold3 prediction and molecular dynamics calculation. Q.Q. designed the fluorescent degron peptide, Y.W. analyzed the chain linkage using mass proteomics. Q.Z. and Y.R. drafted the manuscript. All authors (Y.R., Q.Z., Y.D., Z.X., Q.Q., Y.W., S.C., Y.Y., M.P., L.L) read, discussed and analyzed the manuscript. L.L, M.P., Q.Z. supervised the project.

## STAR**⍰**Methods

### RESOURCE AVAILABILITY

#### Lead Contact

Further information and requests for reagents may be directed and will be fulfilled by Lei Liu, Man Pan or Qingyun Zheng, the correspondence authors.

#### Material Availability

Plasmids and reagents generated in this study will be made available on request, but we may require a payment and/or a completed Materials Transfer Agreement if there is potential for commercial application.

#### Data and Code Availability

Coordinates and cryo-EM maps of Cdc48 complex have been deposited in the Protein Data Bank (PDB) and in the Electron Microscopy Data Bank (EMDB), including Cdc48-Npl4/Ufd1-Ubx2-Ub_n_-Eos (EMD-68128, PDB 21ZW), Cdc48-Npl4/Ufd1-Ubx2-Ub_6_ (EMD-68127, PDB 21ZV).

### EXPERIMENTAL MODEL AND SUBJECT DETAILS

The *Escherichia coli* DH5α strain was used to amplify plasmids and generate mutations. The *Escherichia coli* BL21(DE3) or Rocetta2(DE3) strain were cultured in Luria broth (LB) medium to recombinantly express proteins except Ubr1 and Ufd4. The *Saccharomyces cerevisiae* BY4741 strain was used to recombinantly express yeast Ubr1 and Ufd4.

### METHOD DETAILS

#### Plasmid construction

A soluble construct of full length Faf2 was engineered by replacing the hydrophobic membrane-anchoring region (residues 91-121) with a flexible (GS)_14_ linker. Other variants of Faf2 were generated including FAF2^ΔUBX^ (a C-terminal truncation containing residues 1-357); FAF2^CC-UBX^ (a minimal fragment encompassing the coiled-coil (CC) and UBX domain, residues 275-445); FAF2^ΔCC^ (an internal deletion mutant where residues 275-350 were excised, and the flanking N-terminal and C-terminal segments were tethered directly). The soluble Ubx2 construct was generated by replacing the internal residues from 81 to 172 with a flexible (GS)_14_ linker. Ubx2^CC-UBX^ truncation concludes a minimal fragment encompassing the CC and UBX domain, residues 362-571). All constructs were cloned into pET28a with an N-terminal His-tag for expression in *E. coli* BL21 (DE3).

#### Protein expression and purification

Ubx2, Faf2, and their variants were expressed in Rosetta2 (DE3) cells as His_6_ fusions with an HRV-3C protease cleavage sites. Cells were lysed, clarified by centrifugation, and loaded on a Ni-NTA resin equilibrated with Lysis Buffer (50 mM HEPES, pH 8.0, 500 mM NaCl, 20 mM imidazole and 5 mM MgCl_2_). After washing with 20 CV of Wash Buffer (50 mM HEPES, pH 8.0, 500 mM NaCl, 50 mM imidazole and 5 mM MgCl_2_), HRV 3C protease was added to the resin and incubated at 4 °C overnight to cleave off the tag. Cleaved target proteins were washed off from the resin with SEC Buffer (25 mM HEPES, pH 8.0, 150 mM NaCl and 5 mM MgCl_2_) and further purified by size exclusion chromatography (Superdex 200 increase 10/300).

Wild-type and mutant p97, human Ufd1-Npl4, Ub, Ub^K48R^-Ub, Uba1, gp78RING-Ube2g2, Ubc4 were purified as described^22,18,16,39^. Wild-type and mutant Cdc48, yeast Ufd1-Npl4, Ubc2 and His-SUMO-degron-mEos3.2 (degron-Eos) constructs were purified as described^15,13^. Engineered Ube2e1 and mEos3.2 fused with a C-terminal KEGYEE hexapeptide (Eos-KEGYEE), which serves as a defined acceptor site for enzymatic ubiquitination were purified as described^31^. SFP was purified as described^42^. Ubr1 and Ufd4 were expressed in *S. cerevisiae* and purified by Flag affinity chromatography as described^39,50^.

Briefly, all proteins except Ubr1 and Ufd4 were induced in *E. coli* BL21(DE3) by adding 0.5 mM IPTG when the culture reached an OD_600_ of 0.6-0.8. After incubation at 18°C for 16 h, cells were harvested by centrifugation and resuspended in Lysis Buffer (50 mM HEPES, pH 8.0, 500 mM NaCl, 20 mM imidazole and 5 mM MgCl_2_). To purify the Ufd1-Npl4 complex, the resuspended cells of respective construct were mixed at this step.1 mM PMSF was added to the suspensions, and cells were lysed by sonication. Lysates were cleared by centrifugation for 90 min at 17,000 × *g*. Then the supernatants were flowed through a Ni-NTA gravity column twice at 4 °C. Beads were washed with Wash Buffer (50 mM HEPES, pH 8.0, 500 mM NaCl, 20 mM imidazole and 5 mM MgCl_2_). The His-tagged proteins were eluted in Elution Buffer (50 mM HEPES, pH 8.0, 150 mM NaCl, and 400 mM imidazole). HRV3C protease or SUMO protease (Ulp1) were added to remove the fusion-tag, respectively, and dialyzed in SEC Buffer (25 mM HEPES, pH 8.0, 150 mM NaCl and 5 mM MgCl_2_). Proteins were further polished by size-exclusion chromatography and/or anion-exchange chromatography to ensure high purity. Peak fractions corresponding to the hexameric/monomeric form were pooled and concentrated. The purity of the recombinant proteins was assessed by SDS-PAGE followed by Coomassie Brilliant Blue staining, showing a purity of >95%. Proteins were then aliquoted, flash-frozen in liquid nitrogen, and stored at –80°C.

#### Synthesis of free polyubiquitin chains

K48-linked polyubiquitin chains were assembled by mixing 1 μM Uba1, 10 μM gp78RING-Ube2g2 and 1 mM ubiquitin in the ubiquitination buffer containing 20mM HEPES, pH 7.4, 100mM KCl, 10 mM ATP and 10 mM MgCl_2_ at 37 °C for 4 hours. The reaction mixture was then diluted ten-fold with 50 mM NaOAc, pH 4.5, and separated by a Source S cation exchange column (GE Healthcare). The corresponding peaks were collected and dialyzed in 25 mM HEPES, pH 8.0. Hexa-ubiquitin chain (Ub_6_) was further purified via size-exclusion column equilibrated in the SEC Buffer for cryo-EM studies.

#### Polyubiquitination of degron-substrates

For cryo-EM studies, degron-Eos was directly used for polyubiquitination. For substrate unfolding assays, Eos constructs were first irradiated under long-wave UV lamp for 1 hour at 4 °C to induce the photoconversion of mEos3.2, followed by ubiquitination. The polyubiquitination reaction was performed as previously described^39,13^. Briefly, for K48-polyUb-Eos, 10 μM degron-Eos was mixed with 0.5 μM Uba1, 10 μM Ubc2, 0.3 μM Ubr1 and 0.2 mM ubiquitin in the ubiquitination buffer and incubated at 30 °C for 3 hours.

#### Length-defined ubiquitination of degron-substrates by SUE1 strategy

SUE1-guided free Ub chain transfer reaction was performed as previously described^31^. Briefly, 0.5 μM Uba1, 5 μM Ube2e1, 15 μM different Ub chain with precise length and 10 μM Eos-KEGYEE as substrate in the ubiquitination buffer at 37°C for 2 hours.

#### Preparation of branched ubiquitinated substrates

K29/K48-branched-Eos was generated by grafting K29-linked chains onto pre-assembled K48-linked backbones. Purified K48-polyUb-Eos bearing medium-length Ub chains (10 μM) was mixed with 0.5 μM Uba1, 10 μM Ubc4, 0.5 μM Ufd4 and 0.2 mM ubiquitin in the ubiquitination buffer at 30 °C for 5 hours.

M1/K48-branched-Eos was generated as described. First, an M1 diUb mutant (Ub^K48R^-Ub) was expressed and purified, in which the distal moiety contains a K48R mutation to prevent chain extension on the branch, while the proximal moiety contains a native Ub unit. The assembly reaction was performed by incubating 10 μM degron-Eos with 0.5 μM Uba1, 10 μM Ubc2, 0.3 μM Ubr1 and 0.2 mM Ub^K48R^-Ub in ubiquitination buffer at 30 °C for 3 hours. Ubr1/Ubc2 specifically catalyzes K48-linkages between the C-terminus of the proximal Ub of one molecule and K48 of the proximal Ub of the next, resulting in a K48-linked backbone where each unit carries an M1/K48 branched chain.

#### Assembly of substrate-engaged Cdc48 complex

Cdc48-Ufd1-Npl4 complex was assembled as previously described^5^. A Cdc48 mutant bearing E588Q mutation was used to decrease the heterogeneity of N domains and reduce the unfolding activity of the complex. Two-fold molar excess of Ub_6_ or K48-polyUb-Eos and Ubx2 was added to the Cdc48-yUfd1-yNpl4 complex (or yNpl4^WT^ for K48-polyUb-Eos or yNpl4^A494F^ for Ub_6_), followed by gel filtration using a Superose 6 column (GE Healthcare) equilibrated in the storage buffer. The assembled substrate-engaged Cdc48 complex was flash frozen in liquid nitrogen for structural studies. No nucleotide was supplemented during the assembly.

#### Specimen preparation for single-particle cryo-EM

All samples for cryo-EM were prepared as previously described^18^, with the following modifications. Fluorinated octyl-maltoside (FOM) was used to relieve the preferred orientations. The complex was concentrated to ∼3.0 mg/mL followed by incubation with 2 mM ATP at room temperature for 5 minutes before adding FOM (final concentration 0.0025% v/v). A 3.5-μL aliquot was immediately applied to glow-discharged Quantifoil R1.2/1.3 holey carbon grids (300 mesh). The grids were blotted for 1.5 seconds and plunge-frozen in liquid ethane using a Vitrobot Mark IV (FEI) maintained at 8°C and 100% humidity.

#### Data collection for single-particle cryo-EM and image processing

Cryo-EM data collection for the Cdc48 complexes was performed on a Titan Krios G3i electron microscope (Thermo Fisher Scientific) equipped with Falcon □ and Gatan BioQuantum K2 direct detection cameras operated at 300 kV. Using EPU software (FEI), micrograph movies were automatically acquired at a nominal magnification of 130,000×, corresponding to a calibrated pixel size of 0.932 Å. The total exposure dose was 50 e□/Å², with a defocus range of –1.0 to –2.5 µm.

For the Cdc48-yUfd1-yNpl4-Ubx2-PolyUb-Eos complex, 6237 movies were collected. Beam-induced motion correction and CTF estimation were suffered using MotionCor2^51^ and CTFFIND4^52^ in CryoSPARC^53^. The following data processing was also performed using CryoSPARC. 3,195,086 particles were picked from the datasets and extracted with a box size of 200 pixels (bin 2, 1.864 Å per pixel) using Bob picker. After several rounds of two-dimensional (2D) classification, ab initio reconstruction, and heterogeneous refinement, we selected 701,375 particles and re-extracted to unbin with a box size of 400 pixels (0.932 Å per pixel). Upon NU refinement, we obtained a map with an overall resolution at 2.72 Å. However, the density map of the Ubx2 region was not well determined. These refined particles were then subjected to a round of 3D Classification with ten classes. Two classes were well classified and show density for Ubx2. To further classify the particles, the particles were combined and subjected to NU refinement. Finally, we obtained the map shows Ubx2 density with an overall resolution at 3.09 Å.

For the Cdc48-yUfd1-yNpl4(A494F)-Ubx2-Ub_6_ complex, 6610 movies were collected. After motion correction and CTF estimation, 4,523,649 particles were picked and extracted using Bob picker. After several rounds of 2D classification, ab initio reconstruction along with heterogeneous refinement, the best class containing 968, 051 particleswas subjected to NU refinement, yielding a density map with an overall resolution at 2.78 Å.

These refined particles were also subjected to a round of 3D Classification with ten classes. Three classes were well classified and show density for Ubx2, we merged the three sets of particles and subjected them to NU refinement. Finally, we obtained the map shows Ubx2 density with an overall resolution at 3.20 Å. The Ubx2 density is much better compared to the polyUb-Eos dataset. Finally, by performing local refinement, the cryo-EM map for the Ubx2 obtained at resolutions of 4.08 Å. A composite map of the Cdc48-yUfd1-yNpl4(A494F)-Ubx2-Ub_6_ complex was generated with Chimera^54^.

#### Model building and refinement

The initial models were built using the Cdc48, Npl4 and unfolded ubiquitin portions from the Cdc48-Npl4 complex structure [Protein Data Bank (PDB) ID: 6OA9], and the ubiquitin portion from the ubiquitin structure (PDB ID: 1UBQ) as templates. Model for Ubx2_Ubx_CC-Ufd1_UT3-Cdc48_Ndomain-Ub^prox^_Ub^unfolded^ were generated using the AlphaFold3-prediction. All these model portions were docked into the cryo-EM electron density maps using Chimera^54^, followed by iterative manual adjustments in COOT and real-space refinement using real_space_refine in Phenix^55^. Regions with poor (five Cdc48_N domains) or missing densities (Cdc48_D2 Ring) were not modeled. Structural figures were generated using a combination of Chimera and ChimeraX. Side-chain interactions were analyzed based on AlphaFold3-predicted models to minimize interpretive bias arising from poor side-chain density in the experimental maps. The data collection and structure refinement statistics are provided in table S1.

#### Substrate unfolding assay

Substrate unfolding assay was performed as previously described^16,22,56^. Briefly, Experiments were carried out in Reaction Buffer (50 mM HEPES, pH 8.0, 100 mM KCl, 5 mM MgCl_2_, 0.5 mg/mL BSA, 0.5 mM TCEP and 0.01% Triton) supplemented with an ATP regeneration mixture (5 mM ATP, 30 mM creatine phosphate, and 50 μg/mL creatine phosphokinase). Proteins were pre-incubated in a 96-well plate (Costar, Cat#3686) for 10 min at 37 °C (for p97) or 30 °C (for Cdc48) before adding the ATP regeneration mixture to initiate the reaction. Final concentrations of the reactants varied according to specific experimental setups, with p97/Cdc48-UN-cofactor complexes typically range from 20 to 200 nM (hexamer) and ubiquitinated substrates from 50 to 1000 nM. Fluorescence signal was monitored using a BMG CLARIOstar^®^ plate reader at 540nm excitation and 580nm emission wavelengths with 30-second intervals for 30-60 minutes. Each reaction condition was repeated three to four times. Background fluorescence was measured by mixing the same amount of substrate with 6 M guanidine-HCl and was subtracted from the average of the experimental groups. Normalized fluorescence and the initial velocity of the reaction (first 10 data points corresponding to 5 min of reaction) were plotted and fitted using GraphPad Prism v.8.4.2.

#### Preparation of fluorescently labelled proteins/substrates

For ybbR labeling, Texas Red-CoA conjugate and Sfp enzyme were made as described^42^. The 11-residue ybbR tag was appended to the N terminus of Ufd1. ybbR-tagged proteins were purified similar to wild-type protein and labelled in SFP labelling buffer (50□mM HEPES pH 8.0, 150□mM NaCl, 10□mM MgCl_2_) with stoichiometric amounts of SFP enzyme and a 3-molar excess of Texas Red-CoA conjugate to Ufd1^ybbR^-Npl4 for 1□h in the dark at room temperature. Free dye and SFP were removed using Ni-NTA resin (Smart-Lifesciences, Cat#SA004005). Labelled proteins were exchanged into storage buffer (25□mM HEPES pH 8.0, 150□mM NaCl) using a size-exclusion chromatography (Superdex 200 increase 10/300 column, GE Healthcare), concentrated using a 50 K MWCO centrifugation filter (Amicon), flash frozen in aliquots in liquid nitrogen, and stored at –80□°C.

The BODIPY-labelled polyubiquitination reaction was performed as previously described^13^. Briefly, 10 μM fluorophore-labelled degron peptide (RHGSGC^BODIPY-FL^GAWLLPVSLVKRKTTLAPNTQTASPPSYRALADSLMQ(AEEA-AEEA)K^Biotin^, purchased from GenScript) was mixed with 0.5 μM Uba1, 10 μM Ubc2, 0.3 μM Ubr1 and 0.2 mM Ub in the ubiquitination buffer and incubated at 30 °C for 3 hours. The reaction mixture containing polyubiquinated substrates were incubated with TM Streptactin resin (LABLEAD, Cat#S1180) and washed for four times then eluted by (50 mM HEPES, pH 8.0, 150 mM NaCl, 50 mM D-biotin, 0.5 mM TCEP). The elution was further separated using a Superdex 200 (GE Healthcare) size-exclusion column equilibrated in the SEC buffer and then analyzed using SDS-PAGE, scanned with the corresponding wavelengths and stained with Coomassie blue. Fluorescently labelled substrates modified with K48-polyUb (degron^BODIPY^-K48-Ub_n_, estimated over fifteen ubiquitin subunits) were collected and flash frozen for further studies.

#### AlphaFold prediction

Structural predictions presented were generated in AlphaFolder Server12 (available at https://alphafoldserver.com/). Two specific assemblies were modeled to interrogate species-specific interaction surfaces: (1) The yeast assembly comprised the S. cerevisiae Cdc48 N-domain (residues 22-212), yUfd1 UT3 domain (residues 17-208), the Ubx2 CC-UBX module (residues 362-572), and two ubiquitin segments representing the proximal folded ubiquitin and the C-terminus peptide of unfolded ubiquitin. (2) The human assembly comprised Ufd1 UT3 domain (residues 18-196), FAF2 (specifying residues used, 281-312), and a K48 diub mimic (UbK48C and Ub77C).

#### Fluorescence emission spectra

All fluorescence measurements were carried out on a BMG CLARIOstar^®^ plate reader at 25□°C in SEC buffer with 0.5 mg/mL BSA. Fluorescence spectra were recorded using an excitation wavelength of 480□nm and emission wavelengths from 500-630□nm for FRET pair between BODIPY-FL and Texas Red. FRET between substrate peptide modified with K48-polyUb labelled with BODIPY-FL on degron peptide at residue Cys6 (degron^BODIPY^-K48-Ub_n_) and p97-Ufd1^Texas^^Red^-Npl4 was detected using 5□nM degron^BODIPY^-K48-Ub_n_, 5 nM p97-Ufd1^Texas^^Red^-Npl4 and 5 nM unlabeled p97-UN.

#### FRET-based binding and subunit engagement assays

To dissect the hierarchical assembly and substrate-induced remodeling of the p97 complex, equilibrium titrations were performed in SEC Buffer with 0.5 mg/mL BSA and 2 mM ATPγS. To quantify the affinity for substrate, 1 nM substrate peptide modified with K48-Ub_n_ labelled with BODIPY-FL (degron^BODIPY^-K48-Ub_n_) were titrated with increasing concentrations of p97-Ufd1-Npl4 labeled with Texas Red (Acceptor) on the Ufd1 subunit (p97-Ufd1^Texas^^Red^-Npl4). These titrations were performed in the presence or absence of Faf2 or Faf2 variants. Donor fluorescence was recorded using an excitation wavelength of 480 nm and emission wavelength of 517 nm.

Raw fluorescence intensities were corrected for background fluorescence and dilution factors. A control titration with unlabeled titrant was performed in parallel to define the donor-only fluorescence baseline. The apparent FRET efficiency (*E*) at each titration point was calculated based on donor quenching using equation (1):

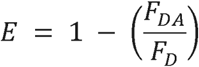

in which *F_DA_* is the donor fluorescence intensity in the presence of the acceptor, and *F_D_*is the donor fluorescence intensity in the presence of the unlabeled control. The concentration dependence of *E* was fitted^57^ to the quadratic binding model (equation 2) to account for ligand depletion effects, as the protein concentrations used were close to the apparent dissociation constant (*K_d_)*:

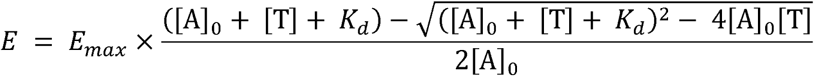

in which [A]_0_ is the total concentration of the fixed fluorescent analyte (Donor), [T] is the total concentration of the titrant (Acceptor), *E_max_* is the FRET efficiency at saturation (representing the maximum extent of subunit engagement or binding), and *K_d_* is the apparent equilibrium dissociation constant. Data fitting and statistical analyses were performed using GraphPad Prism v.8.4.2.

#### Binding Free Energy Calculation (MM/PBSA)

Binding free energies between UT3 and K48 diUb mimic, as well as UT3-K48 diUb mimic in the presence of Faf2, were estimated using the gmx_MMPBSA v1.6.4 tool, which is based on the AMBER MMPBSA.py protocol^45–47^. All calculations were performed using snapshots extracted from 100 ns molecular dynamics trajectories of the respective complexes.

##### Trajectory Preparation

MD trajectories were processed to remove solvent and ions. Each system was centered and wrapped to ensure continuity of the protein-protein complex. Frames were extracted every 10 ps, resulting in 10001 frames per system for analysis.

##### Topology Generation

AMBER topology files for the complex, receptor, and ligand were generated using gmx_MMPBSA from the corresponding GROMACS.tpr and .gro files.

##### MM/PBSA Calculation

The Molecular Mechanics Poisson-Boltzmann Surface Area (MM/PBSA) method was used to compute the binding free energy:

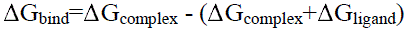

Both electrostatic and van der Waals contributions were included, along with polar solvation energy (Poisson-Boltzmann) and nonpolar solvation energy (LCPO solvent-accessible surface area). Calculations were performed at T = 298.15 K.

## Legends for Supplementary Figures S1-S7

**Figure S1.**
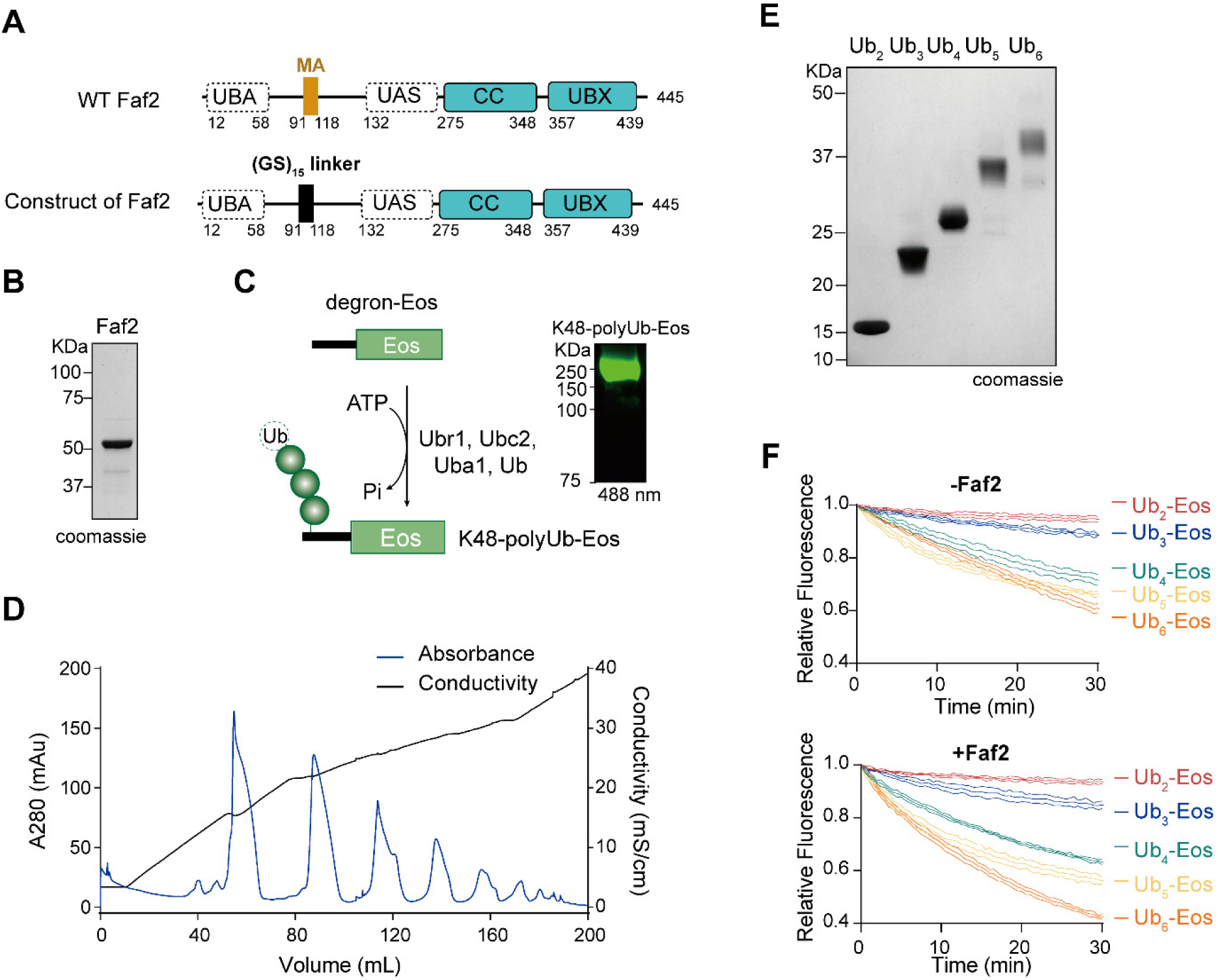
Purification and biochemical characterization of the soluble Faf2 construct and polyubiquitinated substrates. (**A**) Schematic representation of the domain organization of wild-type (WT) Faf2 and the soluble construct used in this study. The hydrophobic membrane anchor (MA, residues 91-118) in WT Faf2 (top) was replaced by a flexible GS_15_-linker to generate the soluble variant (bottom). Key domains are labeled. (**B**) SDS/PAGE analysis of the purified soluble Faf2 construct stained with Coomassie Brilliant Blue. The gel shows a single prominent band corresponding to the recombinant protein. (**C**) Schematic illustration of the enzymatic synthesis of K48-polyubiquitinated Eos (K48-polyUb-Eos) (left). The substrate, comprising Eos fused to a degron, was ubiquitinated in the presence of Uba1 (E1), Ubc2 (E2), Ubr1 (E3), ubiquitin, and ATP. SDS-PAGE analysis of the Eos substrates modified with K48-linked polyubiquitin chains, visualized by fluorescence scanning at 488 nm. (right) (**D**) Purification of K48-linked diUb, triUb, tetraUb, pentaUb and hexaUb by cation-exchange chromatography. The absorbance at 280 nm (blue) reveals distinct peaks eluted by a salt gradient (conductivity, black). (**E**) Fractions corresponding to the peak apexes in (D) were resolved by SDS-PAGE and visualized by Coomassie staining. The migration patterns confirm the isolation of K48-linked ubiquitin chains of defined lengths (Ub_2_ – Ub_6_). (**F**) Representative unfolding traces of K48-Ub_n_-Eos substrates (n = 2 – 6) corresponding to the data in Figure 1F. The fluorescence decay was monitored in the absence (top) or presence (bottom) of Faf2. Note that longer ubiquitin chains trigger faster unfolding, an effect further amplified by Faf2.

**Figure S2.**
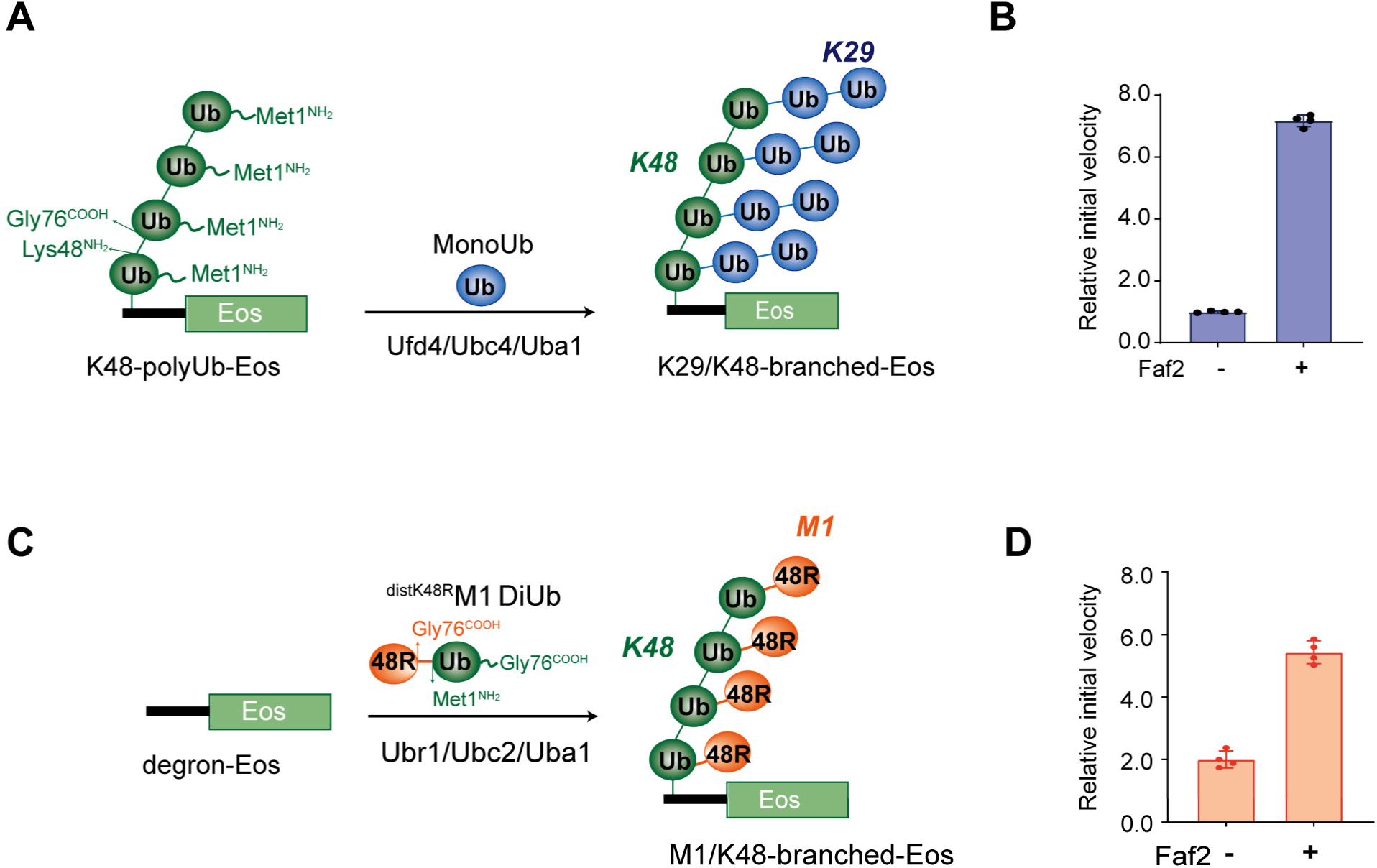
Preparation and unfolding analysis of K29/K48– and M1/K48-branched ubiquitin conjugates. (**A**) Synthesis of K29/K48-branched-Eos. Pre-formed K48-polyUb-Eos was incubated with Uba1, Ubc4, and Ufd4 in the presence of mono-ubiquitin to install K29-linked branches onto the K48-linked backbone. (**B**) Quantification of the unfolding velocities of branched substrates shown in Figure 1G. Relative initial unfolding rates of K29/K48-branched-Eos (blue) in the absence or presence of Faf2. Velocities were normalized to the condition without Faf2. Data represent mean ± s.d. (n = 4 independent experiments). (**C**) Synthesis of M1/K48-branched-Eos. The substrate was generated by Ubr1/Ubc2-mediated polymerization using a M1-linked ubiquitin dimer (Ub^K48R^-Ub) as the building block. This strategy yields a K48-linked backbone where each unit is decorated with a single M1-linked K48R ubiquitin moiety. (**D**) Quantification of the unfolding velocities of branched substrates shown in Figure 1H. Relative initial unfolding rates of M1/K48-branched-Eos (orange). Velocities were normalized to the condition without Faf2. Data represent mean ± s.d. (n = 4 independent experiments).

**Figure S3.**
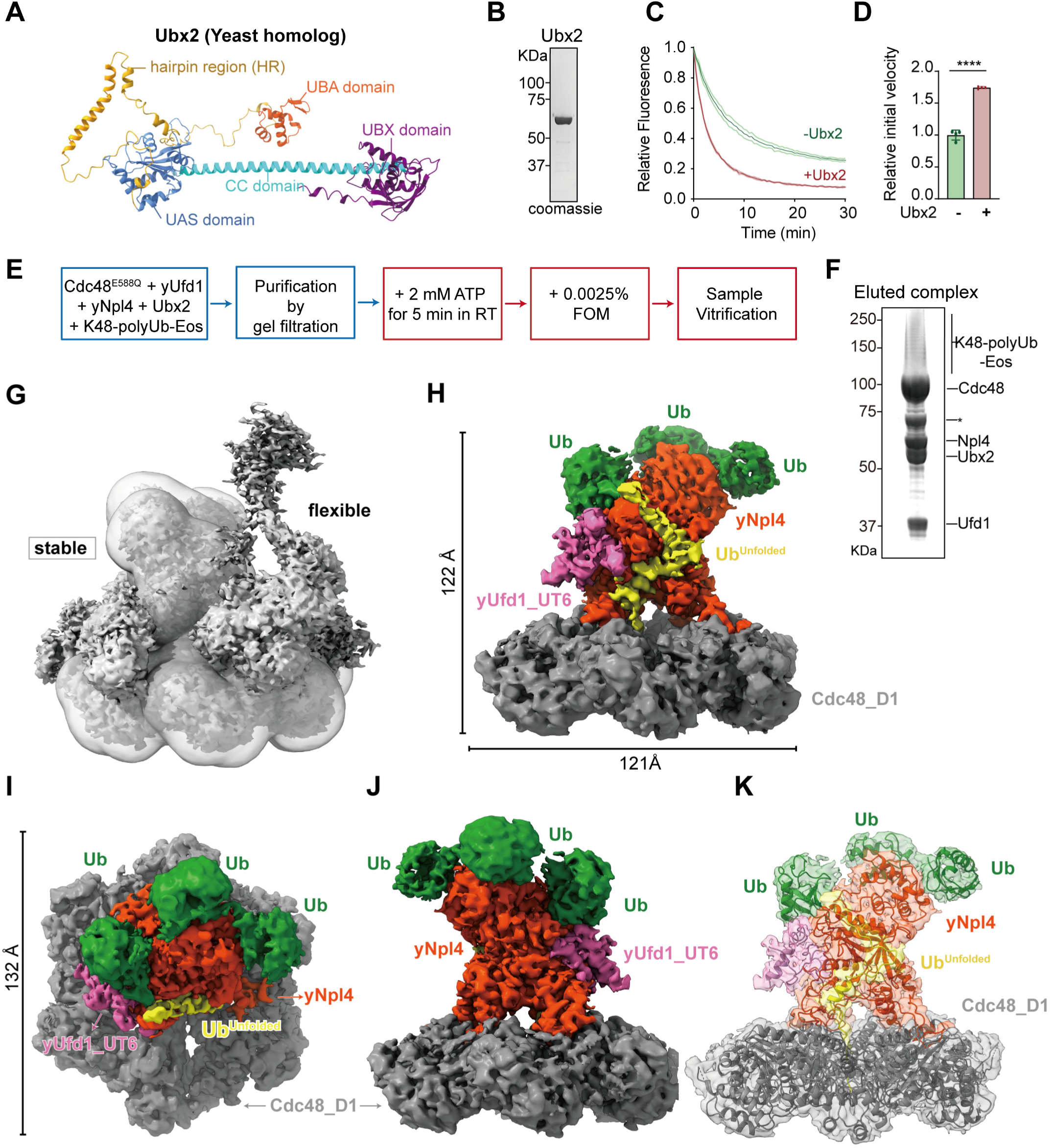
Biochemical characterization and Cryo-EM structure determination of the *S. cerevisiae* Ubx2 and Cdc48-UN complex. (**A**) Structural model of *Saccharomyces cerevisiae* Ubx2 (yeast homolog of Faf2). The predicted structure highlights the N-terminal UBA domain, UAS domain, Coiled-coil region, UBX domain and the hydrophobic hairpin region (HR) which anchors the protein to the membrane. (**B**) Purification of the soluble Ubx2 construct. Coomassie-stained SDS-PAGE showing the purified recombinant Ubx2, in which the hydrophobic hairpin region (residues 81-172) was replaced by a flexible linker to ensure solubility. (**C**) Ubx2 stimulates the unfolding activity of the yeast complex. Representative unfolding traces of K48-polyUb-Eos by Cdc48-Ufd1-Npl4 in the absence (green) or presence (red) of Ubx2. (**D**) Quantification of the initial unfolding velocities from (K). Data represent mean ± s.d. (n = 3). ****, *p* < 0.0001 (two-tailed Student’s *t*-test). (**E**) Sample preparation procedures. Steps performed at 4 °C were in blue boxes. Steps performed at room temperature are in red boxes. (**F**) SDS-PAGE analysis of the purified Cdc48-yUfd1-yNpl4-Ubx2-K48-polyUb-Eos complex. (**G**) The Cryo-EM density map of the Cdc48-yUfd1-yNpl4-Ubx2-K48-polyUb-Eos complex. The central gray transparent region represents the area that can be used for stable structural modeling, while the peripheral portions are highly dynamic and difficult to capture. (**H-J**) Three-dimensional reconstruction of the stable region of the complex by cryo-EM (three different views). (**K**) Atomic model built into the resolvable density of the complex.

**Figure S4.**
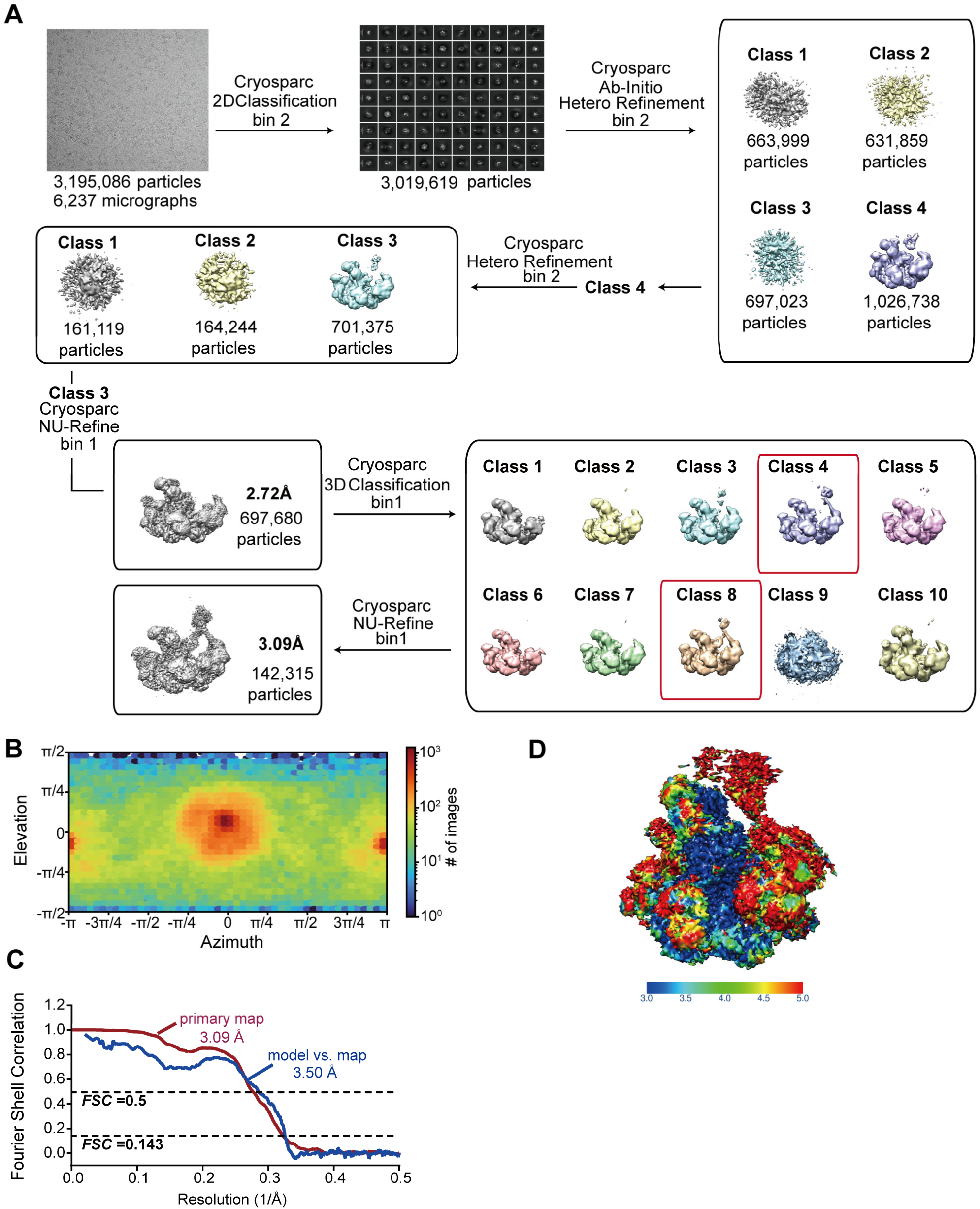
Cryo-EM image processing workflow (Cdc48-yUfd1-yNpl4-Ubx2-K48-polyUb-Eos complex) (**A**) Representative micrograph and flow chart of data processing of the Cdc48-yUfd1-yNpl4-Ubx2-K48-polyUb-Eos complex. (**B**) The Orientational distribution of the particles used for the final reconstructions (from cryoSPARC). (**C**) The gold-standard Fourier shell correlation (FSC) curves from the 3D refinement, the model vs. map FSC curve is also shown. (**D**) Local resolution of primary map calculated using cryoSPARC.

**Figure S5.**
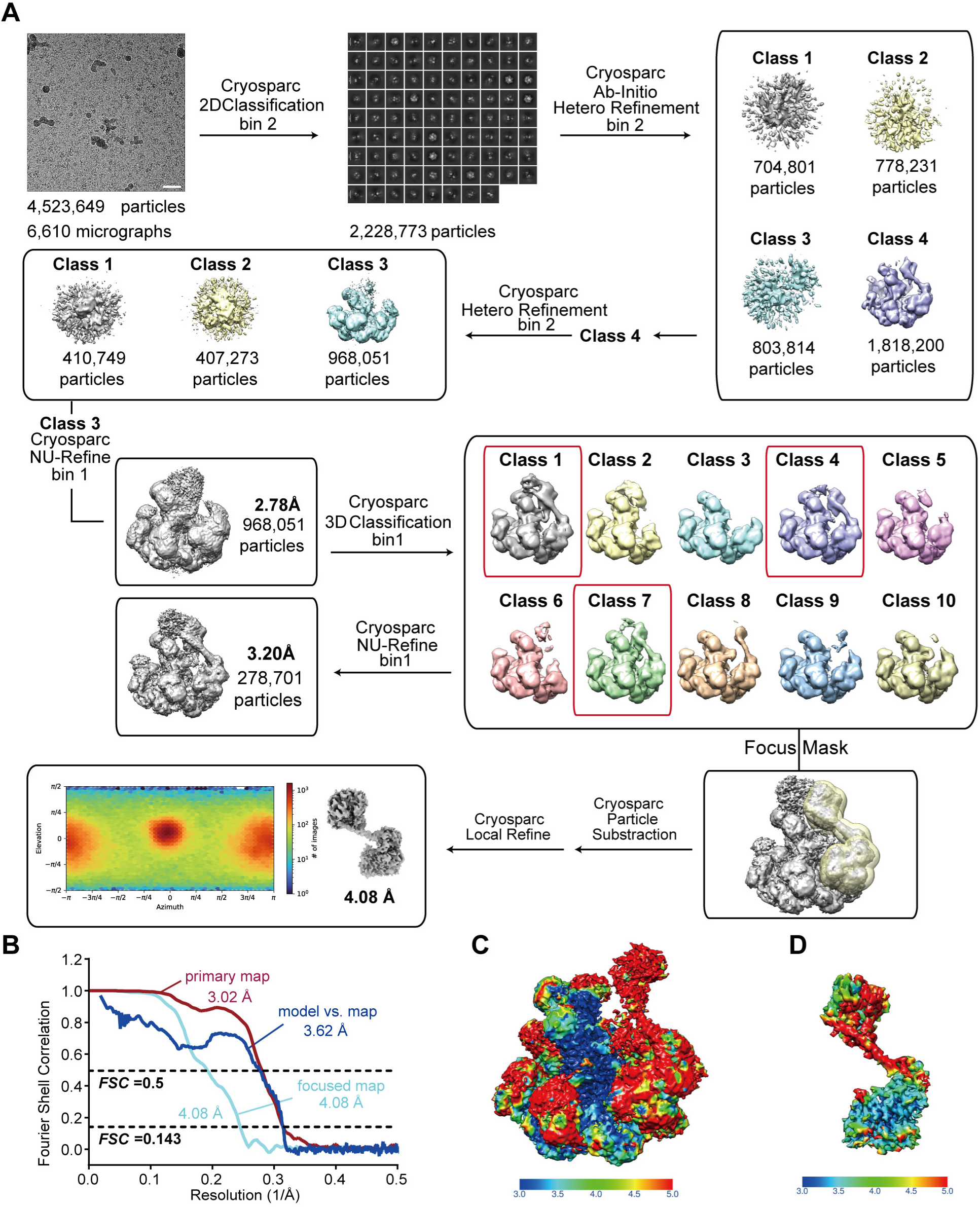
Cryo-EM image processing workflow (Cdc48-yUfd1-yNpl4-Ubx2-K48-Ub_6_ complex) (**A**) Overview of the cryo-EM data processing using cryoSPARC. Masked 3D classifications without realignment were applied to Cdc48^N^-yUfd1^UT3^-Ubx2^CC-UBX^ subcomplex with Ub^prox^ and unfolded Ub^C-term^. Orientational distribution plot demonstrating complete angular sampling. (**B**) The gold-standard Fourier shell correlation (FSC) curve for the overall complex structure and locally refined subcomplex (highlighted in cyan). (**C**) Local resolution of primary map calculated using cryoSPARC. (**D**) Local resolution of focused map calculated using cryoSPARC.

**Figure S6.**
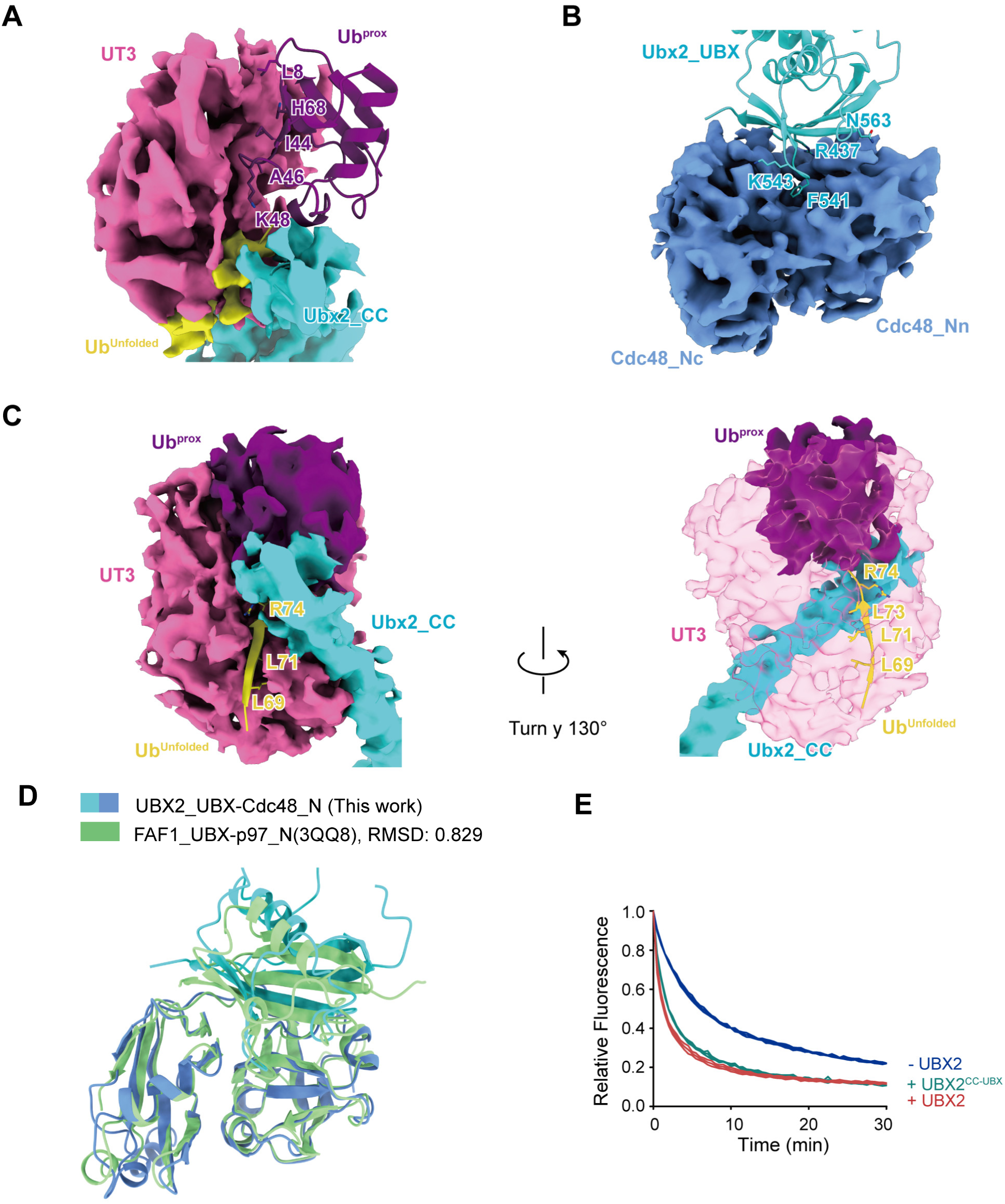
Detailed structural interfaces within the Ubx2-Cdc48-Ufd1-Npl4-substrate complex. (**A**) Recognition of the proximal ubiquitin (Ub^prox^) by yUfd1-UT3. The interaction highlights the binding of the yUfd1-UT3 domain (pink surface) to the canonical I44 hydrophobic patch of the proximal ubiquitin moiety (purple ribbon). Key ubiquitin residues involved in the interface (L8, I44, H68, A46) and the linkage site (K48) are labeled. (**B**) The recruitment interface between the Ubx2 UBX domain and Cdc48. Structural details of the Ubx2 UBX domain (cyan ribbon) docked onto the N-domains of the Cdc48 hexamer (blue surfaces). Labelled residues (R437, F541, K543, N563) delineate the critical contacts mediating the stable anchoring of Ubx2 to the ATPase machinery. (**C**) Interaction between the UT3 domain and the unfolding ubiquitin (Ub^unfolded^). Two rotated views (Turn y 130°) illustrating the contact interface between the UT3 domian (pink surface/transparent) and the ubiquitin moiety positioned for unfolding (yellow ribbon). The interface specifically involves the C-terminal tail of ubiquitin (residues L69, L71, R74), suggesting a role for the CC domain in orienting the substrate for pore insertion. (**D**) A structure alignment of yeast Ubx2^UBX^-Cdc48^N^ determined in this study with Faf1^UBX^-p97^N^ (PDB: 3QQ8). Root-mean-square deviation (RMSD) of backbone atoms are indicated. Faf1^UBX^-p97^N^ is colored in green. (**E**) Unfolding of K48-polyUb-Eos by Cdc48-UN or Cdc48-UN complexed with Ubx2 or Ubx2^CC-UBX^.

**Figure S7.**
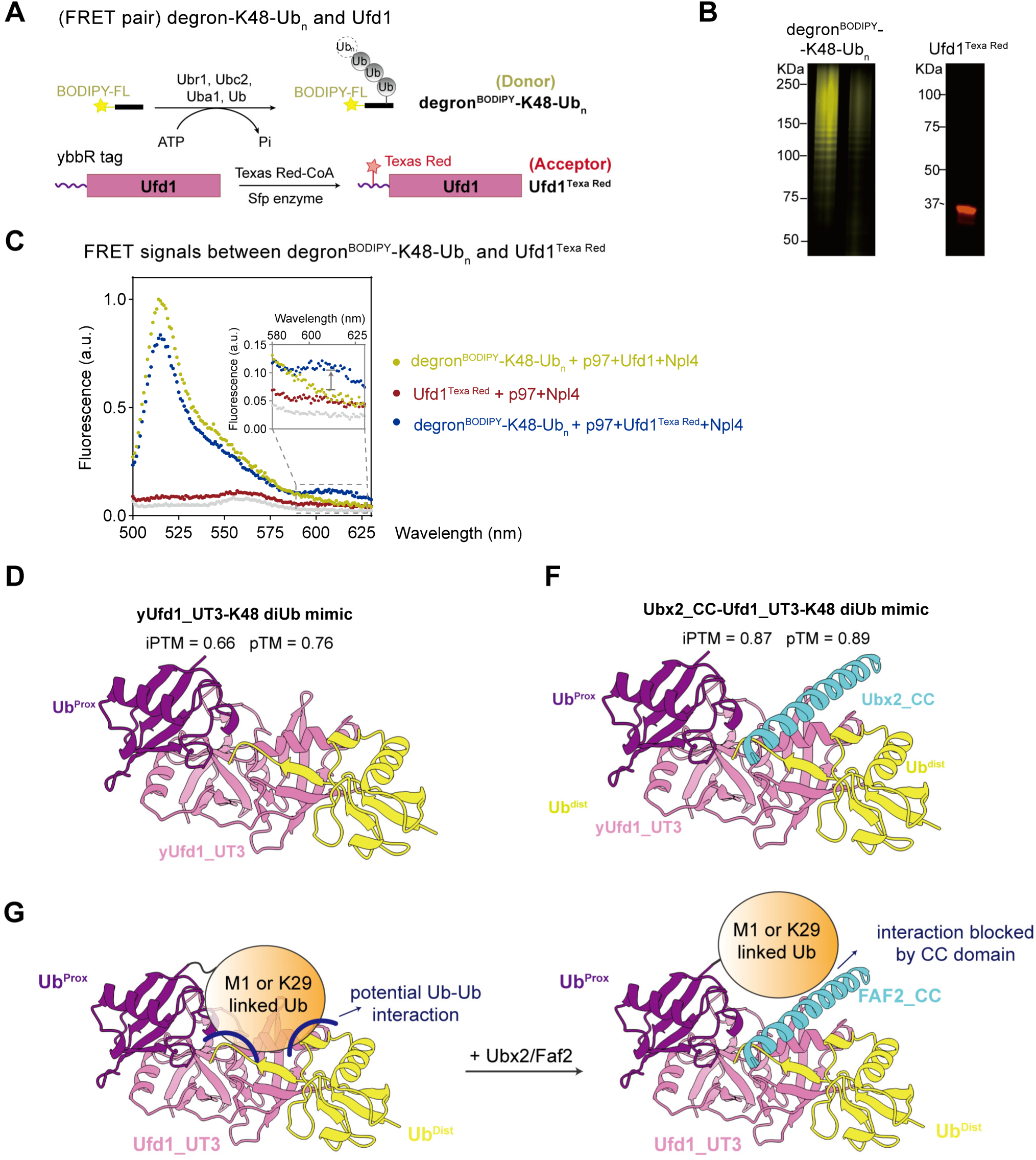
FRET reporter system and structural prediction for functional investigation into CC-UBX domain of FAF2/Ubx2. (**A**) Schematic strategy for generating the FRET pair. Top: The donor probe (degron^BODIPY^-K48-Ub_n_) was generated by enzymatic ubiquitination of a BODIPY-FL labeled degron peptide using the Ubr1 E3 ligase system. Bottom: The acceptor protein (Ufd1^Texas^^Red^) was prepared by site-specific labeling of a ybbR-tagged Ufd1 with Texas Red-CoA using Sfp transferase. (**B**) SDS-PAGE gels visualized under blue light (left) or green light (right) excitation show the characteristic high-molecular-weight smear of the ubiquitinated donor substrate and the single labeled band of the Ufd1 acceptor, respectively. (**C**) Validation of FRET signals. Fluorescence emission spectra (excitation at 480 nm) of p97-Ufd1-Npl4 complexes reconstituted with the indicated probes. Compared to the donor-only control (yellow), the presence of the acceptor (blue) results in quenching of the donor peak (∼515 nm) and the appearance of sensitized acceptor emission (∼610 nm, highlighted in the inset), confirming effective energy transfer. (**D-E**) AlphaFold 3 structural modeling of the yUfd1_UT3-K48-diUb mimic (**D**) and the Ubx2_CC-yUfd1_UT3-K48-diUb mimic (**E**), Predicted structural models and the corresponding confidence scores (iPTM and pTM) generated by AlphaFold 3 are shown. (**F**) Model scheme for Faf2/Ubx2 can confers broadened tolerance to branched K48-polyubiquitin chains. CC domain stabilize the UT3-mediated “open” conformation and blocked the potential Ub-Ub interaction that branched ubiquitination poses to proximal Ub engagement.

**Table S1.**
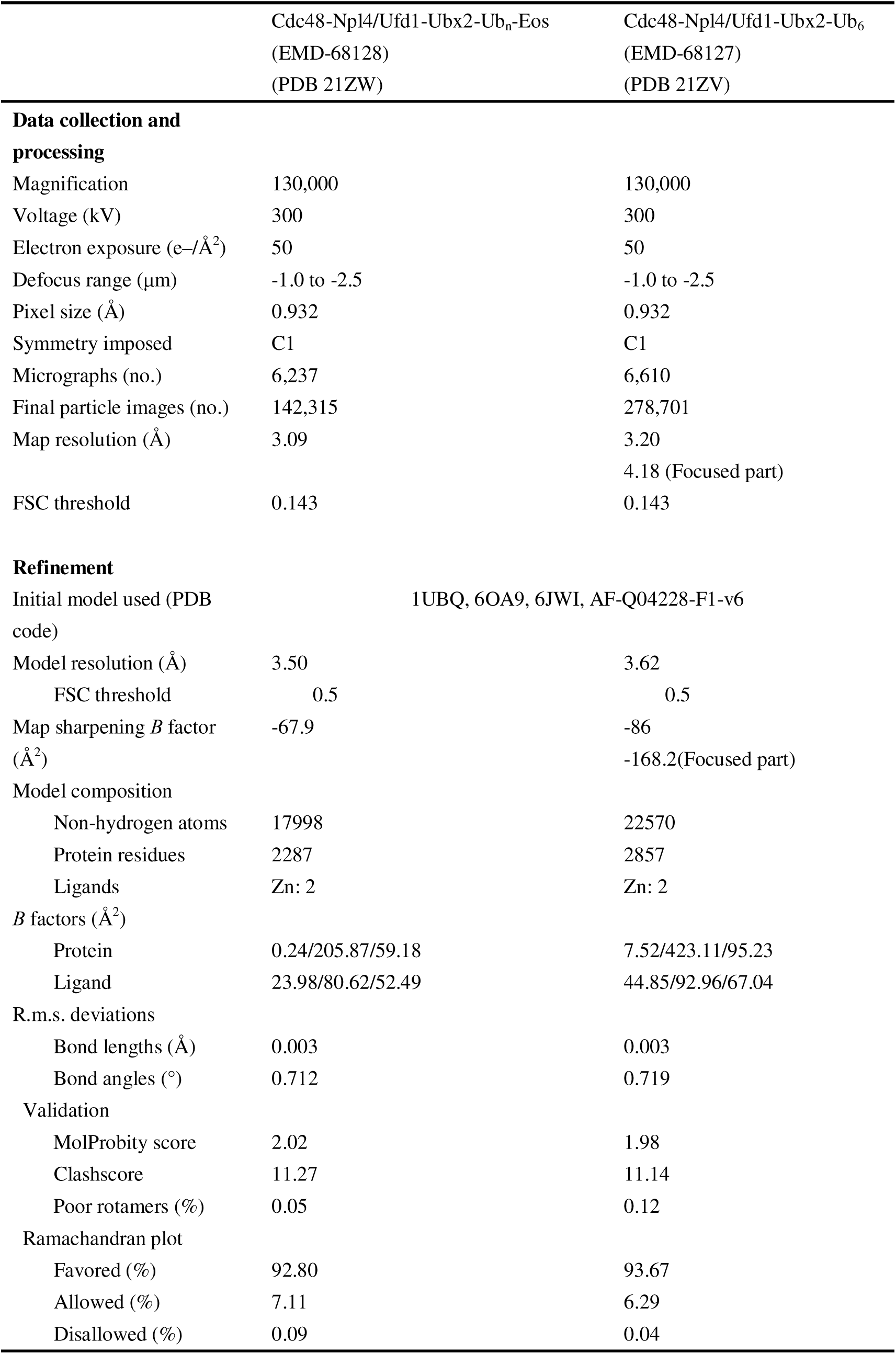
Cryo-EM data collection, refinement, and validation statistics for Ubx2-activated substrate-engaged Cdc48 complex.

**Table S2.**
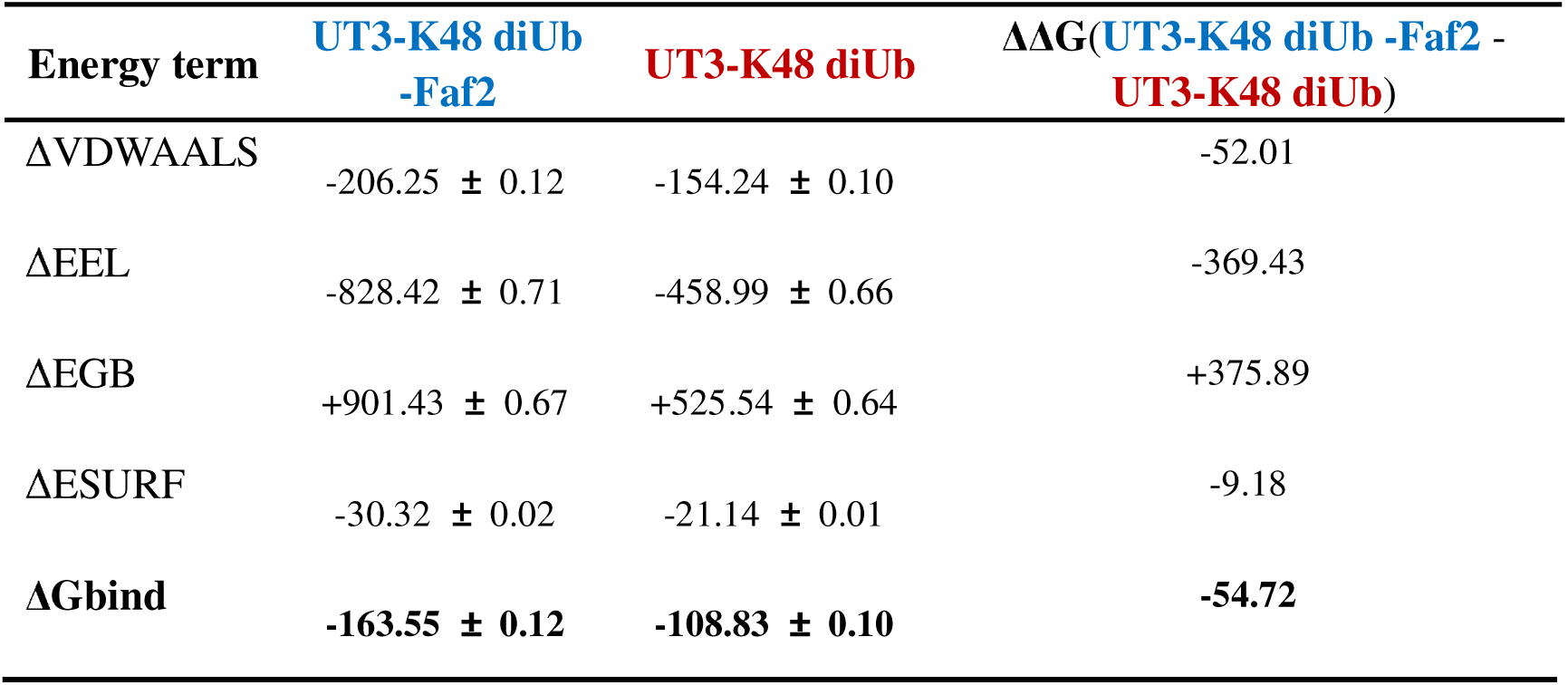
Comparison of MM/GBSA binding free energies for two protein-ligand systems (kcal/mol)

